# Sex chromosome turnover in African annual killifishes of the genus *Nothobranchius*

**DOI:** 10.1101/2024.03.25.586567

**Authors:** Monika Hospodářská, Pablo Mora, Anna Chung Voleníková, Ahmed Al-Rikabi, Sergey A. Simanovsky, Tomáš Pavlica, Marie Altmanová, Karolína Janečková, Jana Štundlová, Nikolas Tolar, Marek Jankásek, Matyáš Hiřman, Thomas Liehr, Martin Reichard, Eugene Yu. Krysanov, Petr Ráb, Christoph Englert, Petr Nguyen, Alexandr Sember

## Abstract

Sex chromosomes of teleost fishes often have low levels of differentiation and undergo frequent turnovers. Annual *Nothobranchius* killifishes comprise representatives with male-heterogametic XY or X_1_X_2_Y sex chromosome systems, scattered across their phylogeny, nested within species lacking cytologically detectable sex chromosomes. They thus provide a suitable system to study sex chromosome evolution and turnover. Here, we combined molecular cytogenetics and genomic analyses to examine several multiple sex chromosome systems in *Nothobranchius* spp. and their outgroup *Fundulosoma thierryi*.

We used fluorescence *in situ* hybridization with three sex chromosome-specific painting probes and bacterial artificial chromosomes (BAC) bearing eight orthologues of genes found to be repeatedly co-opted as master sex determining (MSD) genes in fishes. Our results suggest at least four independent origins of sex chromosomes in the genus *Nothobranchius*. The synteny block carrying *amhr2* gene was shared by X_1_X_2_Y systems of *N. brieni*, *N. guentheri* and *N. lourensi*, but the autosomal additions and the overall neo-Y chromosome structure differed among these species. On the other hand, *gdf6* gene was localized to neo-Y of *F. thierryi*. None of the mapped MSD gene candidates seems to determine sex in *N. ditte*.

We further sequenced genomes of *F. thierryi* female and *N. guentheri* male by long-read platforms and performed analyses of male and female Pool-seq data and coverage to delimit their non-recombining regions, determine degree of their differentiation, and thus complement the cytogenetic data in assessing potential MSD genes. We found low level of sex chromosomes differentiation in *F. thierryi.* In *N. guentheri*, however, we identified two distinct evolutionary strata on neo-Y. The *amhr2* gene resides in the younger stratum and has low allelic variation, which questions its role in sex determination.

## Introduction

Sex chromosomes represent one of the most intriguing topics of contemporary evolutionary biology. These specialized chromosomes usually develop from an ordinary autosomal pair through acquisition of a primary signal in the sex determination pathway (Charlesworth et al. 2005; Wright et al. 2016; Ponnikas et al. 2018). Around the newly established sex-determining region (mostly on Y in male or on W in female heterogametic systems, respectively), the recombination is usually arrested or greatly reduced, which allows for the independent evolution of X and Y or Z and W chromosomes. As the non-recombining region spreads, the Y/W chromosome degenerates by various mechanisms and eventually becomes cytologically distinctive (i. e. heteromorphic) with respect to its X/Z gametolog (recently reviewed in Kratochvíl et al. 2021). The pace of differentiation and degeneration is not always well correlated with the evolutionary age of a given sex chromosome system (Zhou et al. 2014; Charlesworth 2021; Kuhl et al. 2021). One of the processes, which can restart sex chromosome differentiation, is sex chromosome turnover, i. e. replacement of ancestral sex chromosome system by a new one (Saunders 2019; Vicoso 2019; Meisel 2020).

Despite more than a century of intense research (Abbott et al. 2017), the field of sex chromosome evolution continues to capture the attention of researchers as many pressing questions still wait to be satisfactorily answered. Among these are i) primary mechanism of recombination suppression between sex chromosomes, and ii) drivers of sex chromosome turnover and its role in reproductive isolation and adaptation (Kratochvíl et al. 2021; Olito et al. 2022; Smith et al. 2023; Kitano et al. 2024).

While most mammals and birds have stable and highly differentiated sex chromosomes (Cortez et al. 2014; Zhou et al. 2014), teleost fishes posses remarkably variable sex chromosome systems, with frequent turnovers and often low genetic differentiation (Volff et al. 2007; Kitano & Peichel 2012; Gamble 2016; Schartl et al. 2016; Sember et al. 2021). Different sex chromosome systems at various stages of differentiation (or convergent evolution of the same system from non-homologous autosomes) may be found within closely related species and even at the level of distinct conspecific populations (Kitano & Peichel 2012; Cioffi et al. 2017; El Taher et al. 2021; Sember et al. 2021; Kitano et al. 2024). Besides that, fishes have the highest known diversity of sex-determining (MSD) genes among animals, evolved independently in non-related species by various molecular mechanisms (Herpin & Schartl 2010; Guiguen et al. 2018; Kitano et al. 2024). Many of these so-called “usual suspects” belong to TGF-ß pathway, namely *gsdf* (gonadal soma derived factor), *gdf6* (growth differentiation factor), *amh* (anti-Müllerian hormone) and its receptor *amhr2* (Pan et al. 2021; Kitano et al. 2024). Thanks to all the above mentioned features, teleost fishes provide vital models for sex chromosome research.

African annual killifish genus *Nothobranchius* Peters, 1868 (Cyprinidontiformes: Nothobranchiidae) occupies extremely variable environment of ephemeral water bodies in savannahs of southeastern Africa. They are small-bodied, highly sexually dimorphic, and dichromatic freshwater fishes, with larger and more colourful males (Wildekamp 2004). Because their natural lifespan is constrained by drying out at the begining of dry season, *Nothobranchius* spp. have evolved key adaptations such as exceptionally short life span with accelerated growth and sexual maturation, and desiccation-resistant diapause embryos (Furness 2016; Cellerino et al. 2016; Vrtílek et al. 2018). Given these properties, *Nothobranchius* spp., and particularly the turquoise killifish *N. furzeri*, a vertebrate with the shortest natural lifespan in captivity, became a suitable model for a variety of biomedical research, particularly for research on aging (Cellerino et al. 2016; Hu & Brunet 2018).

The genus *Nothobranchius* currently comprises more than 90 species (Nagy & Watters 2022; Fricke et al. 2024) placed in seven main evolutionary clades (van der Merwe et al. 2021). All species live in small, geographically isolated, populations but may occasionally disperse and colonize new water ponds during major floods in rainy season. Genetic drift processes thus considerably affected their genome evolution and diversification (Bartáková et al. 2013; Cui et al. 2019; van der Merwe et al. 2021). According to van der Merwe et al. (2021), the estimated age of the genus *Nothobranchius* is 13 million years (MY). Available data suggest relatively recent diversification within the clades (Dorn et al. 2014; van der Merwe et al. 2021), implying fast rate of ongoing allopatric speciation (Bartáková et al. 2013; Dorn et al. 2014).

As for genome organization, *Nothobranchius* killifishes underwent frequent karyotype changes in majority of the phylogenetic clades, with an important contribution of fusions and fissions, leading to a wide range of diploid chromosome numbers (2n = 16–50) and striking variability in karyotype composition found in 73 representatives examined to date (Krysanov & Demidova 2018; Krysanov et al. 2016, 2023). Genomes of *Nothobranchius* spp. contain high amounts of repetitive DNA (Reichwald et al. 2009, 2015; Cui et al. 2019; Štundlová et al. 2022), which may facilitate chromosome changes (Štundlová et al. 2022; Lukšíková et al. 2023; Voleníková et al. 2023).

Regarding sex chromosomes, only male-heterogametic systems have been identified thus far in *Nothobranchius*. Two sister species, *N. furzeri* and *N. kadleci*, share homeologous ♀XX/♂XY sex chromosome system (Reichwald et al. 2015; Štundlová et al. 2022), while six other representatives possess multiple sex chromosome system of the ♀X_1_X_1_X_2_X_2_/♂X_1_X_2_Y type which is also present in the outgroup species *Fundulosoma thierryi* (Ewulonu et al. 1985; Krysanov et al. 2016; Krysanov & Demidova 2018). This system has most probably resulted from Y-autosome fusion, hence the X_1_ denotes the original X chromosome and X_2_ represents the unfused homolog of the chromosome which has been added to Y chromosome. While available data is not sufficient to tell whether these multiple sex chromosome systems have a single origin or evolved independently, the latter hypothesis is favoured (Krysanov & Demidova 2018) as the species in question, namely *N. brieni*, *N. ditte*, *N. guentheri*, *N. janpapi*, *N. lourensi*, and *Nothobranchius* sp. Kasenga, have different morphology of sex chromosomes and are scattered across the *Nothobranchius* phylogeny, nested within species with unknown sex chromosome constitution (Krysanov & Demidova 2018; van der Merwe et al. 2021; Bartakova et al. in rev.).

A candidate MSD gene has been identified thus far only in *N. furzeri* and *N. kadleci* where multiple lines of evidence (Reichwald et al. 2015; Štundlová et al. 2022) including recent functional analyses (Richter et al. 2023) strongly suggest that Y-linked allele of *gdf6* gene (*gdf6Y*) determines the male sex in these species. *Gdf6Y* evolved via allelic diversification in the original locus and differentiation of the X and Y sex chromosomes displays high degree of inter-population polymorphisms (Reichwald et al. 2015; Štundlová et al. 2022). We previously hypothesized that the recombination rate between these XY sex chromosomes could be reduced due to heterochiasmy (i.e., a sex-specific pattern of recombination) (Štundlová et al. 2022).

Degree of differentiation of X_1_X_2_Y sex chromosomes in *Nothobranchius* has been so far assessed only indirectly, based on constitutive heterochromatin distribution and patterns of mapped repetitive sequences (Lukšíková et al. 2023; Voleníková et al. 2023). With the sole exception of *N. brieni*, these analyses showed generally low degree of accumulation of heterochromatin and repetitive sequences along with minor differences between gametologs, corroborating the standard patterns found in majority of fish multiple sex chromosomes (Sember et al. 2021).

In the present study, we combined molecular cytogenetic and genomic methods to decipher whether *Nothobranchius* X_1_X_2_Y sex chromosome systems had recurrent and independent origins, and to assess their differentiation and potential MSD genes. We revealed at least four sex chromosome turnovers in *Nothobranchius* killifishes and *F. thierryi*, with the two prime candidates for MSD role from TGF-ß pathway, *gdf6* and *amhr2*, being present in the neo-Y-linked non-recombining region, however, in the latter not in the oldest evolutionary stratum in *N. guentheri*.

## Material and Methods

### Fish species sampling

We analyzed four *Nothobranchius* species with X_1_X_2_Y sex chromosome system, namely *N. brieni*, *N. guentheri*, *N. ditte* and *N. lourensi*, and the outgroup species *Fundulosoma thierryi* carrying the same type of multiple sex chromosomes. To compare the results to our earlier findings (Štundlová et al. 2022), we also included *N. furzeri* with the XY sex chromosome system. A single population per species has been sampled except for *N. guentheri* (two populations). The studied individuals from *N. furzeri* were sampled from laboratory population (see Bartáková et al. 2015; Blažek et al. 2017). The remaining species were obtained from specialists and experienced hobby breeders who keep strictly population-specific lineages derived from original imports. In this case, the species identity was confirmed based on key morphological characters (Wildekamp 1996, 2004; Nagy 2018). The sampling is summarized in Table 1.

**Table 1.**
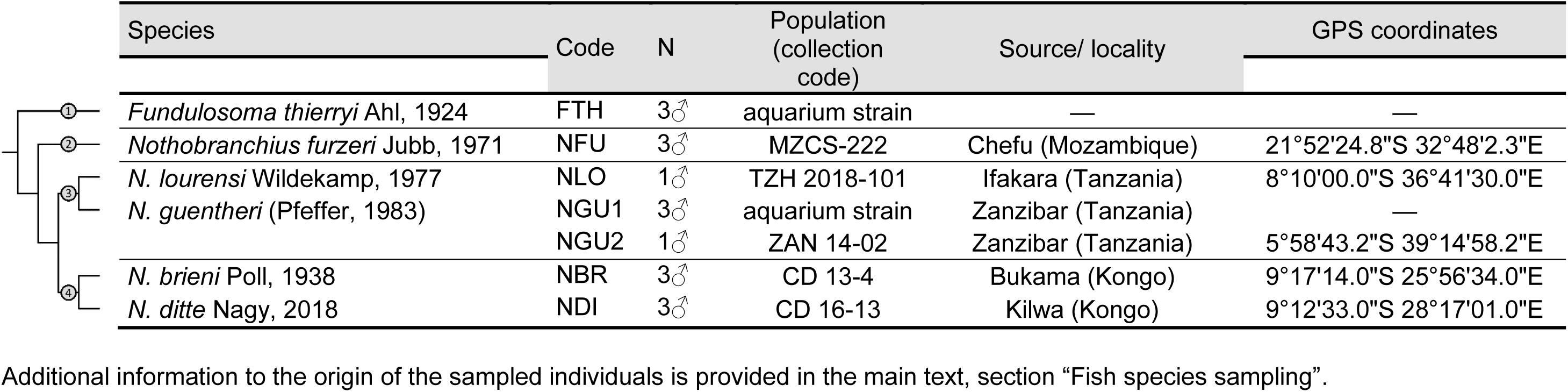
List of *Nothobranchius* killifish species used in this study, with assignment to their phylogeographic lineage: 1) outgroup, 2) Southern, 3) Coastal and 4) Kalahari clade, population/collection codes, sample sizes (N) for cytogenetic analyses, source/geographic origin and GPS coordinates of sampling localities. Phylogenetic relationships follow **van der Merwe et al. (2021**) and the exact placement of *N. lourensi* in the phylogeny is based on recent phylogenomic analysis of Tanzanian *Nothobranchius* species using double-digest restriction site-associated DNA (ddRAD) sequencing (**Bartakova & Reichard 2023**).

### Conventional cytogenetics

Mitotic chromosome spreads were obtained either from regenerating caudal fin tissue or from cephalic kidney. In the former, we followed Völker & Ráb (2015), with the modifications by Sember et al. (2015), and with an altered time of fin regeneration (one-to-two weeks; in *N. lourensi* up to three weeks). For the kidney-derived chromosome preparations, we followed either Ráb & Roth (1988), or Kligerman & Bloom (1977). The latter protocol was modified according to Krysanov & Demidova (2018). The chromosomal spreading quality was enhanced following Bertollo et al. (2015). Preparations were inspected under phase-contrast for the amount and quality of chromosome spreads. Suitable slides were dehydrated in an ethanol series (70%, 80% and 96%, 2 min each) and stored in −20 °C.

### DNA isolation

High-weight molecular genomic DNA (HMW gDNA) from male individuals of all species under study, and from females of *F. thierryi* and *N. guentheri*, for the purpose cytogenetic experiments and pool-seq, was extracted from liver, spleen and brain tissue using a MagAttract HMW DNA Kit (Qiagen, Hilden, Germany) following the manufacturer’s instructions. However, volume of reagents and buffers were doubled except for the elution buffer. Genomic DNA (gDNA) was used for molecular cytogenetic applications and short-read genome sequencing. HMW gDNA for long-read sequencing was extracted using the Nanobind tissue kit (PacBio, Menlo Park, CA, USA) following the application note for isolation of HMW DNA from frozen tilapia testis tissue.

### Whole chromosome painting (WCP) and Zoo-FISH (cross-species fluorescence *in situ* hybridization) with sex chromosome-specific probes

To assess the degree of shared synteny between multiple sex chromosomes and to verify the mechanism of their origin, we prepared three WCP probes generated from multiple sex chromosomes of distinct *Nothobranchius* species. In the case of *N. guentheri* and *N. lourensi,* we cut out easily discernible large neo-Y chromosome from mitotic metaphases (10 and 8 copies, respectively). In the case of *Fundulosoma thierryi*, we did not have suitable mitotic slides for this procedure, therefore we isolated a whole sex-trivalent from meiotic spreads (8 copies), based on its characteristic morphology (see Supplementary Fig. 1a).

The glass needle-based microdissection was carried out under inverted microscope (Zeiss Axiovert 135, Zeiss, Oberkochen, Germany) using sterile glass needle attached to a mechanical micromanipulator (Zeiss) (a procedure detailed in Al-Rikabi et al. 2020). The primary chromosome material was amplified and labelled (by SpectrumOrange-dUTP or SpectrumGreen-dUTP; both Vysis Downers Grove, IL, USA) in two subsequent reactions of degenerate oligonucleotide-primed (DOP) PCR following Yang et al. (2009). The resulting painting probes were designated as NGU-Y (neo-Y chromosome of *N. guentheri*), NLO-Y (neo-Y of *N. lourensi*), and FTH-triv (entire sex trivalent of *F. thierryi*). The final probe cocktail for each slide was composed of two compared WCP probes (100 ng each) supplemented with 9 μg of unlabeled blocking DNA, to block non-specific hybridization by highly repeated DNA sequences (4.5 μg DNA of an analyzed species and the remaining 4.5 μg comprising equal ammount of gDNA of species from which the WCP probes have been generated). Zoo-FISH procedure was performed using a combination of two previously published protocols. Specifically, the slide pre-treatment, probe/chromosome denaturation and hybridization followed Sember et al. (2015), while post-hybridization washing was done according to Yano et al. (2017) with slight modifications described in Štundlová et al. (2022). After standard post-hybridization washes, the slides were passed through an ethanol series and mounted in antifade containing 1.5 µg/mL DAPI (4′,6-diamidino-2-phenylindole; Cambio, Cambridge, United Kingdom).

### Fluorescence *in situ* hybridization (FISH) with 5S and 18S rDNA probes

Our previous results from Chromomycin A_3_ staining (Voleníková et al. 2023) suggested that the additional WCP signals (see Results section and Supplementary Fig. 2), could colocalize with rDNA clusters. To test it, we used FISH with 5S and 18S rDNA fragments. For the probe preparation, we used previously cloned PCR fragments of 18S rDNA from *N. guentheri*, and 5S rDNA from *N. kadleci*, both successfully used in our previous study (Štundlová et al. 2022). The probe composition and FISH conditions followed Sember et al. (2015) with slight modification specified in Štundlová et al. (2022).

### Bacterial arifical chromosome (BAC) clone isolation and nick-translation

We mapped *N. furzeri* BAC clones (Table 3) carrying genes commonly involved in fish sex determination onto chromosomes of the studied species. The BAC clones were selected by BLASTn search using *N. furzeri* BAC end sequences in its genome sequence, selecting those clones for which both tags map within 100 Kb of candidate genes (Reichwald et al. 2015) (Table 3). BAC DNA was isolated using NucleoBond Xtra Midi kit (Macherey Nagel, Düren, Germany) and verified by PCR with specific primers (Supplementary Table 1). Labelling of isolated BAC DNA was done by the nick-translation method. The reaction contained 2 µg of BAC DNA, 1× nick translation buffer (5 mM Tris-HCl, 0.5 mM MgCl_2_, 0.0005% bovine serum albumin, BSA; pH 7.5), 10 mM β-mercaptoethanol, 50 µM dATP, 50 µM dCTP, 50 µM dGTP, 10 µM dTTP, 20 µM Cy3-dUTP or Fluorescein-12-dUTP, 40 U DNA polymerase I (Thermo Fisher Scientific, Waltham, MA, USA) and 0.01 U DNase I (RNase-free, Thermo Fisher Scientific). The reaction was incubated at 15 °C for 5 h and finally stopped by adding a 1× loading buffer (50% glycerol, 250 mM EDTA, 5.9 mM bromphenol blue).

### BAC-FISH

The FISH experiments were done using a combination of three previously published protocols (Sember et al. 2015; Yano et al. 2017 and Sahara et al. 1999), with slight modifications described in Štundlová et al. (2022). Briefly, the hybridization mixture was composed of 300 ng of Cy3-labeled BAC DNA and 500 ng of FITC-labeled BAC DNA, 6 µg of thermally fragmented male competitive gDNA (99 °C, 20 min), 1500 ng of DOP-PCR-derived competitive DNA from male gDNA (based on Yang et al. 2009) and 25 µg of sonicated salmon sperm DNA (Sigma-Aldrich, St. Louis, MO, USA). The hybridization mixture was denatured for 5 min at 90 °C. After application of the probe cocktail on the slide, the hybridization took place in a moist chamber at 37 °C for 3 days. Subsequently, non-specific hybridization was removed by post-hybridization washes. Specifically, slides were treated once 5 min at 62 °C in 1% Triton X-100 in 0.1× SSC and then 2 min at room temperature in 1% Triton X-100 in 2× SSC. Final wash 2 min at room temperature in 1% Kodak PhotoFlo in H_2_O was followed by counterstaining with 0.2 g/mL DAPI mounted in antifade based on DABCO (1,4-diazabicyclo(2.2.2)-octane; Sigma-Aldrich). Slides reused for repeated FISH experiments were reprobed following Yoshido et al. (2015).

In some instances, WCP and BAC-FISH probes were successfully mapped simultaneously in a combined experiment, following the WCP protocol, as described above.

### Microscopic analyses and image processing

Images from majority of cytogenetic experiments were captured under immersion objective 100× using BX53 Olympus microscope (Olympus, Tokyo, Japan) equipped with appropriate fluorescence filter set, and coupled with a black and white CCD camera (DP30W Olympus). Images were acquired for each fluorescent dye separately using DP Manager imaging software (Olympus). The same software was used to superimpose the digital images with the pseudocolours (blue for DAPI, red for Cy3 and SpectrumOrange, green for FITC or SpectrumGreen). Composite images were optimized and arranged using Adobe Photoshop, version CS6 (Adobe Systems, San Jose, CA, USA). Images from BAC-FISH and most of the combined WCP/BAC-FISH experiments were captured using Leica DM6 B Fluorescence Microscope (Pragolab, Prag, Czech Republic) equipped with Leica sCMOS Monochrome Camera DFC9000GT, using the Leica Application Suite X (LAS X) imaging software v3.7.3.23245.

At least 20 chromosome spreads per individual in each method were analyzed, some of them sequentially. The description of chromosome morphology, based on Levan et al. (1964), was modified as: m – metacentric, sm – submetacentric (biarmed chromosomes), st – subtelocentric, and a – acrocentric (monoarmed chromosomes).

### Read control quality and genome size estimation using GenomeScope

To assemble their genome sequence, we sequenced *F. thierryi* male genome and *N. guentheri* female genome using Nanopore (Oxford Nanopore Technologies, Oxford, UK) and PacBio (Pacific Biosciences, Menlo Park, CA, USA) technology, respectively. We also re-sequenced male and female gDNA using Illumina (Illumina, San Diego, CA, USA) to correct long reads and perform coverage analyses.

The raw Illumina sequencing data underwent quality assessment using FastQC (Andrews et al. 2010) and were subsequently trimmed with Trimmomatic v0.39 (Bolger et al. 2014). Oxford Nanopore Technology (ONT) and Pacific Biosciences (PacBio) reads were quality-checked using the NanoPack tool (De Coster et al. 2018), specifically utilizing NanoPlot for assessment and NanoFilt for filtering and trimming. Based on the visualized quality of the sequencing, ONT reads under 10 kb in length and with a quality score below 9 were excluded from the dataset. The Ratatosk tool (Holley et al. 2021) was employed with default settings to correct ONT reads using the accurate Illumina reads from the same sample. In case of PacBio data, only reads longer than 5 kb were used for genome assembly.

For estimating genome size, GenomeScope2 (Ranallo-Benavidez et al. 2020) was used with the Jellyfish k-mer counting tool (Marçais & Kingsford 2011). Various k-mer lengths ranging from 15 to 45 were used, with k=45 ultimately selected for its optimal model fit value for *F. thierryi* and 21 for *N. guentheri*.

### Genome de novo assembly and its quality evaluation

The *F. thierryi* long reads were first corrected using the Illumina short reads. Then, several options were used for genome assembly using Flye v2.8 (Kolmogorov et al. 2019) with the “– –nano-raw” and “*--min-overlap 5000*“ option giving the best assembly in terms of contiguity and BUSCO (Benchmarking Universal Single-Copy Orthologs) completeness. Moreover, an appropriate genome size estimated by GenomeScope2 was also set in the assembly process. The assembly underwent one round of long read polishing via medaka (https://github.com/nanoporetech/medaka) followed by a round of short read polishing using NextPolish (Hu et al. 2020) with Illumina reads. Haplotypic duplicates were then removed using the purge_dups v1.0.1 tool (Guan et al. 2020). Assembly quality was assessed using BUSCO v5 with the Actinopterygii database (odb_10) (Manni et al. 2021). Contamination checks were performed using BlobTools v1.0 (Laetsch & Blaxter 2017), and any contigs associated with non-target organisms were eliminated using the “*seqfilter*” function.

The best result for *N. guentheri* assembly was achieved using Flye v2.8 (Kolmogorov et al. 2019) with the “––pacbio-raw” option, genome size not defined, and without further polishing or de-duplication. Assembly quality was verified with BUSCO v5 with the Cyprinodontiformes database (odb_10) and QUAST v4 (Gurevich et al. 2013).

### Genome annotation

Functional and structural annotations were conducted using the GenSAS v6.0 pipeline (Humann et al. 2019). Repetitive sequences were identified employing RepeatModeler2 (Flynn et al. 2020) utilizing the RMBlast search engine and modules including TFR v4.09, RECON, and RepeatScout v1.0.5. Additionally, TAREAN (Novák et al. 2017) was utilized for satDNA annotation. All consensus sequences marked as satellites by TAREAN, regardless of confidence levels, were integrated into a custom database as dimers to enhance the satellite DNA annotation. RepeatMasker v4.1.1 (Smit et al. 2013–2015, available at http://www.repeatmasker.org) with the NCBI/RMBlast search engine was employed for repeat annotation. This involved a combination of the newly identified repeats from RepeatModeler2 and the custom database containing the satDNA sequences from TAREAN.

RNA-seq data were used for *N. guentheri* genome annotation. To that end, total RNA was extracted from brain tissue dissected from *N. guentheri*. Subsequently, the 150 bp Illumina reads were aligned to the respective genomes using STAR v2.7.7 (Dobin & Gingeras 2015). The genome index required for mapping was generated using the following command: *STAR --runThreadN 9 --runMode genomeGenerate --genomeDir ./genomedir -- genomeFastaFiles <genome> --genomeSAindexNbases 13.* Following index generation, mapping was executed using the command: *STAR --runThreadN 9 --genomeDir ./genomedir --readFilesIn ./Forward.fq ./Reverse.fq*. The resulting SAM file was converted to BAM format using the SAMtools suite (v1.11) (Li et al. 2009). The generated BAM file was then utilized for gene prediction through BRAKER2 with default settings, which incorporates Augustus and GeneMark-EP (Lomsadze et al. 2014; Brůna et al. 2021). For the annotation of *Fundulosoma thierryi* genome assembly, Augustus v3.3.1 (Stanke & Morgenstern 2005) and GeneMark-ES were used directly, without RNA-Seq evidence to guide the process. Furthermore, tRNA and rRNA sequences were identified in all assemblies using tRNAscan-SE v2.0.7 (Chan & Lowe 2019) and RNAmmer v1.2 (Lagesen et al. 2007), respectively.

### Reference guided scaffolding

To obtain pseudochromosome level assemblies of both, *F. thierryi* and *N. guentheri*, we aligned genome assembly contigs to chromosomes of the *N. furzeri* reference (GenBank acc. no. GCA_014300015.1) using Minimap2 v2.17 (Li 2018) and sorted and oriented the contigs into pseudomolecules using RaGOO (Alonge et al. 2019) with “*-b -s*” options.

### Pool-seq analysis

To identify male-specific (MSY) regions, we generated pooled samples of genomic DNA from *N. guentheri* (28 males, 29 females) and *F. thierryi* (20 males, 11 females) and sequenced them in paired-end mode on the Illumina platform. The Illumina pools were quality checked with FastQC v0.11.5 (Andrews et al. 2010) and filtered with the “*--nextseq-trim=20 --minimum-length=100*” options using cutadapt v1.15 (Martin 2011) and trimmed with Trimmomatic v0.36 (Bolger et al. 2014) with following parameters: “*SLIDINGWINDOW:4:25 MINLEN:100 HEADCROP:4 CROP:140*”. The trimmed and filtered reads from female and male pools were mapped separately to the reference genomes using BWA-MEM v0.7.17 (Li & Durbin 2009) with default parameters. Using Picard Toolkit (https://broadinstitute.github.io/picard/) we sorted resulting bam files by coordinate (“*SortSam SORT_ORDER=coordinate*”) and removed PCR duplicates (“*MarkDuplicates REMOVE_DUPLICATES=true REMOVE_SEQUENCING_DUPLICATES=true*”). Subsequently, we generated a file with the nucleotide composition of all genomic positions using the *pileup* function of the software Pooled Sequencing Analysis for Sex Signal (PSASS; Feron & Jaron 2021). We used PSASS to identify non-overlapping 50 kb windows enriched in sex-specific SNPs, using the following parameters: “*--min-depth 10, --freq-het 0.5 --range-het 0.15 --freq-hom 1 --range-hom 0.05 -- window-size 50000 --output-resolution 50000 --group-snps*”.

### Coverage analysis

The short Illumina reads from three male and female samples of both *F. thierryi* and *N. guentheri* were quality checked with FastQC v0.11.5 (Andrews et al. 2010) and filtered with “*- -nextseq-trim=20 --minimum-length=100*” options using cutadapt v1.15 (Martin 2011) and trimmed with Trimmomatic v0.36 (Bolger et al. 2014) with following parameters: “*SLIDINGWINDOW:4:25 MINLEN:100 HEADCROP:10 CROP:130*”. Repetitive sequences pseudochromosome-level assemblies were identified by RepeatModeler v1.0.11 (Smit et al. 2008–2015, available at http://www.repeatmasker.org) and annotated by RepeatMasker v4.0.7 (Smit et al. 2013–2015, available at http://www.repeatmasker.org) with the NCBI search. Filtered reads were mapped to the corresponding masked reference via Bowtie2 v2.2.9 (Langmead & Salzberg 2012) with the “*--very-sensitive-local --no-discordant --no-mixed*” parameters and the outputs were compressed to BAM format using SAMtools *view* (v1.3.1; Li et al. 2009) and merged according to sex using SAMtools *merge*. The resulting BAM files were then parsed using utilities from the Bedtools suite v2.25.0 (Quinlan & Hall 2010). A genome file was parsed from the BAM files using SAMtools *view* and divided into 50 kbp sliding windows using Bedtools *makewindows* with “*-w 50000 –s 50000*” parameters. The merged BAM files were sorted with SAMtools *sort* and converted to BED format using Bedtools *bamtobed* “*-split*”. Finally, the per base coverage of aligned sequences within 50 kbp windows spanning the genome was computed using Bedtools *coverage*. In both sexes, coverage depths for each scaffold were normalized by mean coverage across scaffolds and compared between sexes, formulated as the Log2 of the male:female (M:F) coverage ratio. The resulting data, together with those from Pool-seq analysis, were visualised using “*SexGenomicsToolkit/sgt*r” R package (https://github.com/SexGenomicsToolkit/sgtr).

## Results

### Genome assemblies

The raw read counts and lengths for genome sequencing samples are detailed in Supplementary Table 2. A summary of genome assembly statistics for both species is outlined in Table 2. The assemblies exhibited contig N50 values of 6 Mbp and 15 Mbp for *F. thierryi* and *N. guentheri*, respectively. BUSCO analyses showed about 90 % of conserved orthologs to be complete and unique across both species (Table 2). Consequently, both assemblies serve as high-quality references for downstream analyses.

**Table 2.** Summary statistics for *F. thierryi* and *N. guentheri* genome assemblies.

| Assembly summary | *Nothobranchius guentheri* | *Fundulosoma thierryi* |
| --- | --- | --- |
| Total length (Mbp) | 940 | 983 |
| N fragments | 1,021 | 1,621 |
| N50 (Kbp) | 15,275.3 | 6,062.3 |
| Largest contig (Mbp) | 54.174 | 28.078 |
| BUSCO (C; D; F; M) (%) | 90.7; 0.9; 0.7; 8.6 | 90; 1; 0.6; 8.4 |

BUSCO = Benchmarking Universal Single-Copy Orthologs (C = completed; D = duplicated; F = fragmented; M = missing)

**Table 3.**
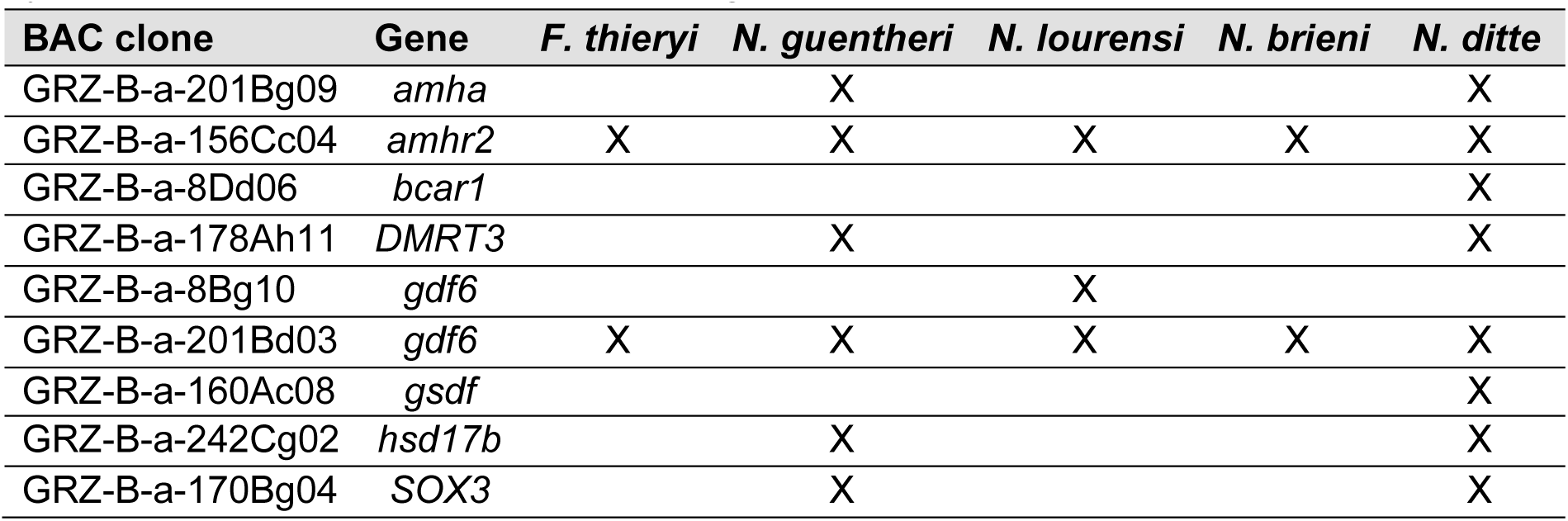
BAC clones from *N. furzeri* genome library which carry specific genes (following information from supplement S1I in Reichwald et al. 2015) used in fluorescence *in situ* hybridization. BAC clones used in screening of particular species are marked as “X”.

| BAC clone | Gene | *F. thieryi* | *N. guentheri* | *N. lourensi* | *N. brieni* | *N. ditte* |
| --- | --- | --- | --- | --- | --- | --- |
| GRZ-B-a-201Bg09 | *amha* |  | X |  |  | X |
| GRZ-B-a-156Cc04 | *amhr2* | X | X | X | X | X |
| GRZ-B-a-8Dd06 | *bcar1* |  |  |  |  | X |
| GRZ-B-a-178Ah11 | *DMRT3* |  | X |  |  | X |
| GRZ-B-a-8Bg10 | *gdf6* |  |  | X |  |  |
| GRZ-B-a-201Bd03 | *gdf6* | X | X | X | X | X |
| GRZ-B-a-160Ac08 | *gsdf* |  |  |  |  | X |
| GRZ-B-a-242Cg02 | *hsd17b* |  | X |  |  | X |
| GRZ-B-a-170Bg04 | *SOX3* |  | X |  |  | X |

### Pool-seq and Coverage analysis

Pool-seq data of both *F. thierryi* and *N. guentheri* were mapped to the reference chromosome level assembly of *N. furzeri* to identify sex-linked regions. Pool-seq analyses identifed regions of increased Fst and male-specific SNPs on chromosomes 3 and 14 in *F. thierryi* (Fig. 1) and chromosome 11 and 13 in *N. guentheri* (Fig. 2). Pool-seq data analyses were complemented by coverage analyses comparing sequencing depth between sexes (Figs 1 and 2), which can identify well-differentiated sex-linked sequences. While the coverage was the same across all chromosomes in *F. thierryi*, the analysis identified well differentiated region on chr. 13 in *N. guentheri* (Fig. 2), clearly revealing two strata of different age in the non-recombining region.

**Fig. 1.**
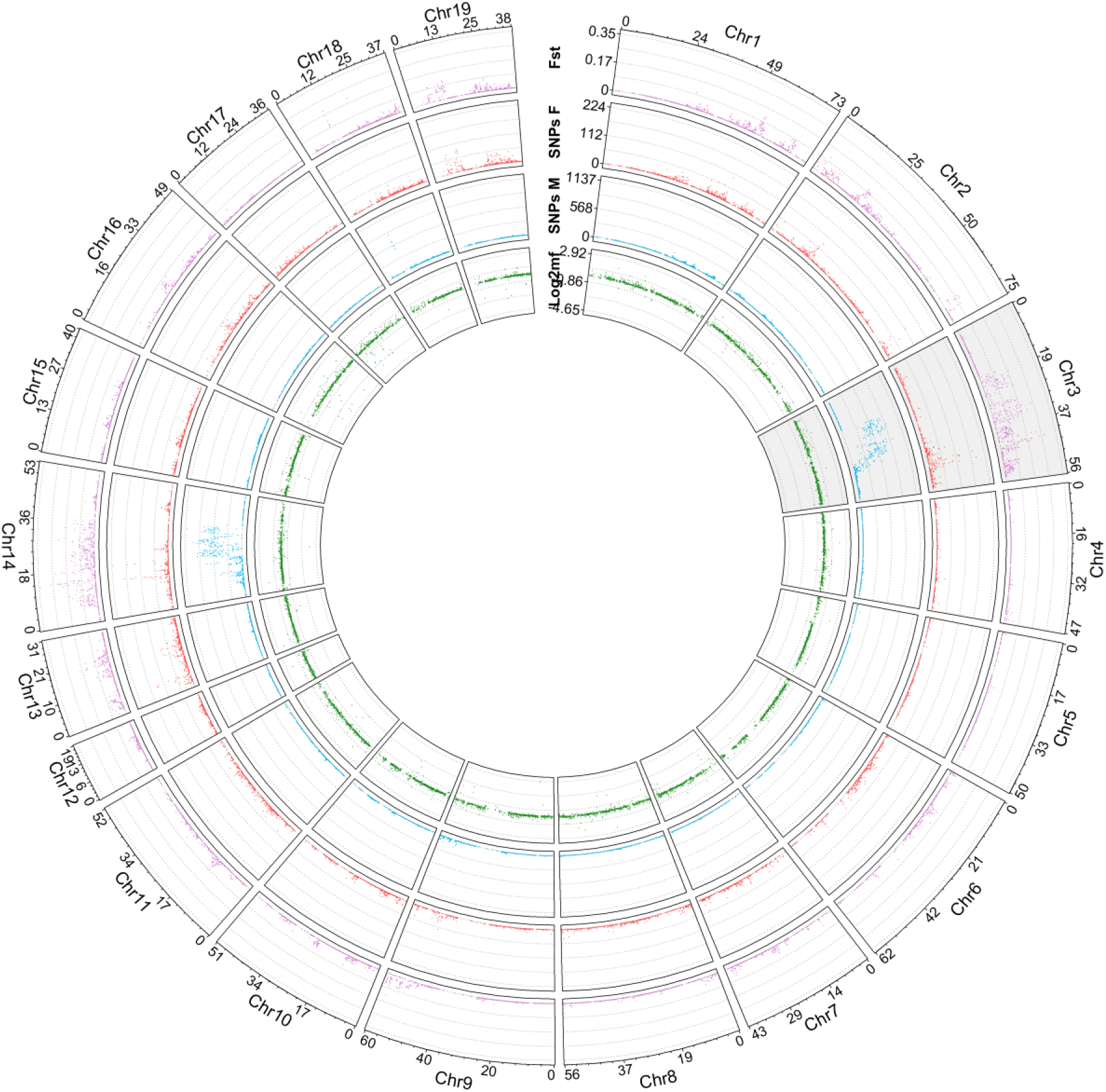
Identification of sex-linked regions in *F. thierryi*. Circo plots show results of coverage (green) and Pool-seq analyses (male and female specific SNPs and relative divergence (Fst) are in blue, red and violet, respectively) using pseudochromosomes constructed with *N. furzeri* as a reference. The sex chromosome of *N. furzeri* is shaded in gray.

**Fig. 2.**
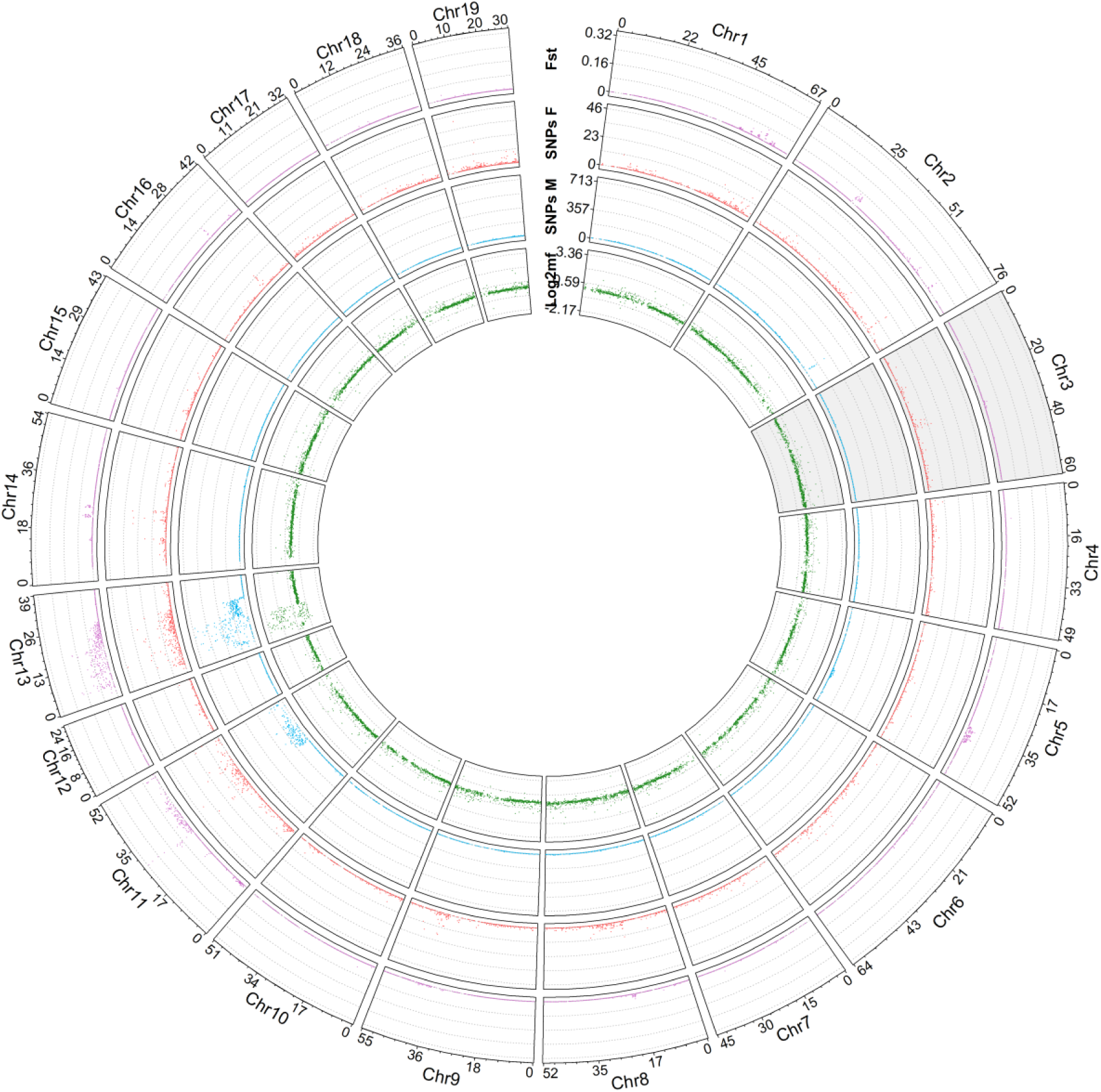
Identification of sex-linked regions in *N. guentheri*. Circo plots show results of coverage (green) and Pool-seq analyses (male and female specific SNPs and relative divergence (Fst) are in blue, red and violet, respectively) using pseudochromosomes constructed with *N. furzeri* as a reference. The sex chromosome of *N. furzeri* is shaded in gray.

### BAC-FISH

We managed BAC screening of potentially sex-determining genes across the sampled species (Supplementary Fig. 2). In outgroup species, *F*. *thierryi*, we identified *gdf6* as the putative MSD gene, similarly as was reported in *N*. *furzeri* and *N*. *kadleci* (Reichwald et al. 2015; Štundlová et al. 2022; Richter et al. 2023). This also corresponds to results observed from pool-seq analysis, which revealed that multiple sex chromosomes of *F*. *thierryi* corresponds to chromosome 14 and chromosome 3 (sex chromosome) of *N*. *furzeri* (Fig. 1). Thus, they share the master sex determining gene *gdf6*, located on chromosome 3. On the contrary, in all other tested species, BAC clone with the *gdf6* gene hybridized to autosomes. In both species from the Coastal clade, *N*. *guentheri* and *N*. *lourensi*, only the gene *amhr2* hybridized to sex chromosomes and thus seems to be a putative MSD gene in this particular clade. Also here, our results from pool-seq analysis in *N*. *guentheri* correspond with localization of *gdf6* on autosomes and not on sex chromosomes. Similarly, in *N*. *brieni* we identified *amhr2* as putative MSD gene. However, any of the tested ”usual suspects” hybridized to sex chromosomes of *N*. *ditte*. The data hence point on the presence of at least two MSD genes in the Kalahari clade: *amhr2* and one another so far unknown.

### Basic chromosomal characteristics

All individuals and analysed populations have been already subject of our previous cytogenetic studies (Štundlová et al. 2022; Lukšíková et al. 2023; Voleníková et al. 2023), in which we confirmed the previously reported diploid chromosome numbers (2n) and the presence of sex chromosomes (Krysanov & Demidova 2018).

### Revese-FISH WCP experiments

The identity of all three sex chromosome-specific painting probes was confirmed by their hybridization back against the chromosomal background of males of respective species. Each probe painted all three members of X_1_X_2_Y system in *F. thierryi*, *N. guentheri* and *N. lourensi*, from end to end (Supplementary Fig. 2).

### Detection of chromosomal homeologies by Zoo-FISH experiments

All hybridization patterns are provided in Supplementary Fig. 2 and schematized in Fig. 3. Reverse FISH experiments (i.e. against the metaphases of the original species) confirmed the specifity of generated WCP probes by marking all members of the X_1_X_2_Y system from end to end, revealing thereby unambiguously the identity of all sex chromosomes in the complement. NGU-Y probe produced additional signals on varied number of autosomes in different species due to the presence of major ribosomal DNA (rDNA) clusters on neo-Y and X_1_ chromosome, as was revealed by rDNA FISH experiments (see Supplementary Fig. 1).

**Fig. 3.**
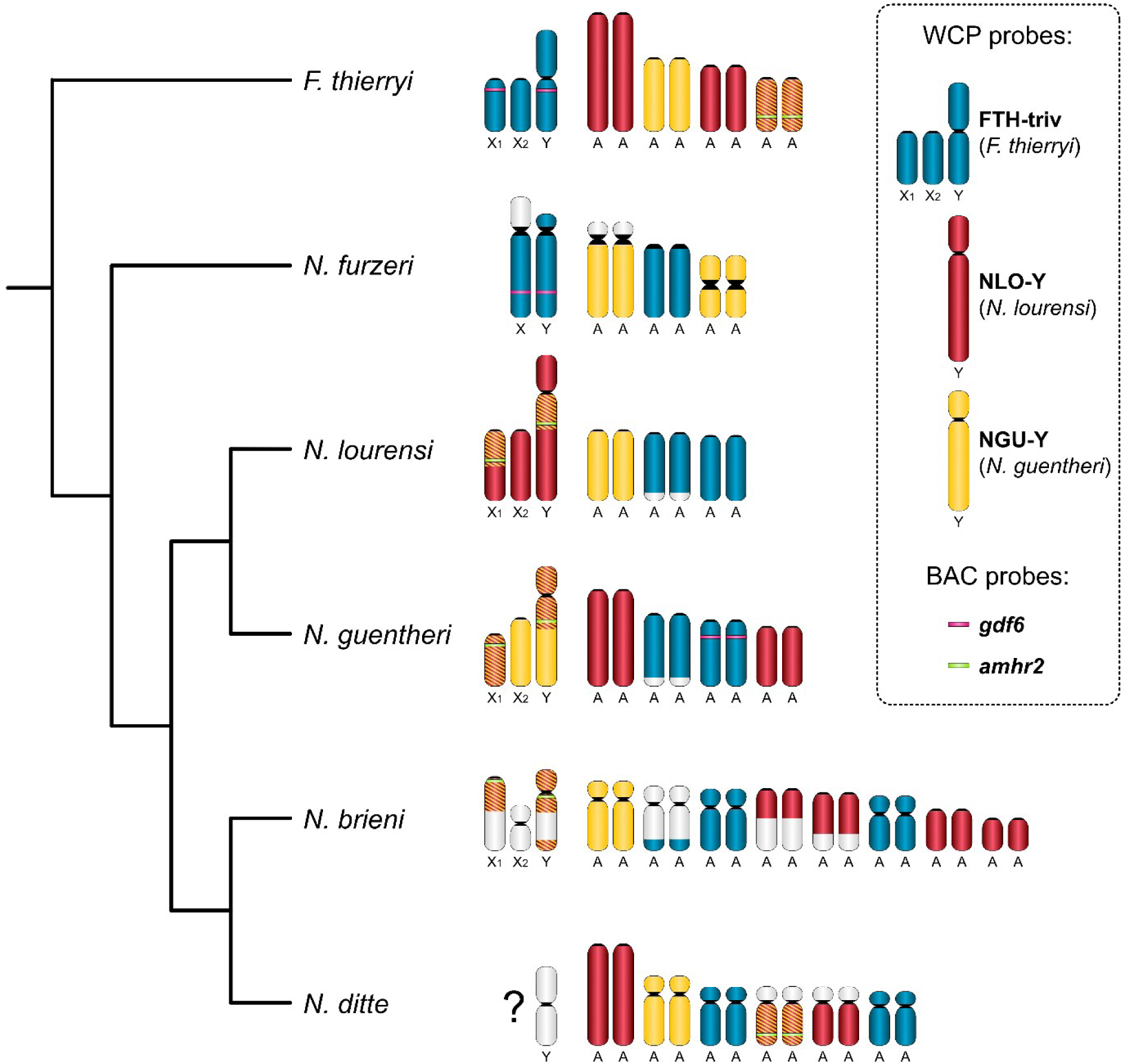
Schematic representation of WCP and BAC-FISH hybridization patterns. The content and colour coding of WCP and BAC-FISH probes is depicted on the right panel. FTH-triv = meiotic sex trivalent from *F. thierryi*; NLO-Y = neo-Y chromosome from *N. lourensi*; NGU-Y = neo-Y chromosome from *N. guentheri*. Segments in crosshatched red-yellow colour represent overlap between NGU-Y and NLO-Y probe. *Gdf6* and *amhr2* = candidate genes for sex-determining role contained in the mapped BAC-clones; their approximate location is depicted on sex chromosomes and, where unambiguously assigned to certain chromosome pair, also on autosomes. A = autosomes. A truncated phylogenic tree follows van der Merwe et al. (2021) and for *N. lourensi* Bartakova et al. (in rev.). Note that none of the WCP probes hybridized to X_1_X_2_Y sex chromosomes of *N. ditte*.

Concerning cross-species (Zoo-FISH) experiments, in *F. thierryi*, each painting probe stained different chromosomes but NGU-Y and NLO-Y entirely overlapped with their signal pattern on a single pair of small acrocentric chromosomes. While NGU-Y probe painted four chromosomes, NLO-Y stained six of them.

In *N. furzeri*, FTH-triv and NGU-Y probes both produced signals on four chromosomes, with non-overlapping distribution patterns. The probes, however, left large (peri)centromeric regions unstained, due to different repetitive DNA content, as may be inferred from our previous study (Voleníková et al. 2023). The FTH-triv probe painted a pair of small acrocentric chromosomes and the long arms of XY sex chromosomes, which were clearly heteromorphic, in line with our previous report (Štundlová et al. 2022).

In *N. lourensi*, the FTH-triv probe entirely painted four acrocentric autosomes. NGU-Y probe marked, besides a pair of another acrocentric autosomes, also distinct segments of two sex chromosomes, where the signal overlapped with NLO-Y probe. Specifically, the two probes co-localized in the segment placed proximally on the long arms, covering about one fourth of the neo-Y length. Complementary experiments by BAC-FISH showed that this segment contains *amhr2* gene. The homologous segment was placed proximally on a medium sized acrocentric sex chromosome, covering the proximal half of this chromosome. Given the presence of this segment, the sex chromosome was denoted as X_1_.

The signal patterns in *N. guentheri* were as follows: the FTH-triv probe hybridized entirely to four acrocentric autosomes. The NLO-Y probe co-localized with the NGU-Y probe in the proximal half of the neo-Y chromosome and over the entire length of the smaller one from the two acrocentric X chromosomes. Given its content (together with the confirmed presence of *amhr2* locus by BAC-FISH), this chromosome was denoted as X_1_. The location of *amhr2* on neo-Y chromosome was very close to the fusion breakpoint. This region could be sometimes apparent as an unlabelled constriction on the long neo-Y chromosome arms (Supplementary Fig. 2), which is consistent with the presence of rDNA clusters in this region (see Supplementary Fig. 1C, D).

Regarding *N. brieni*, the most informative was the NGU-Y probe which, besides staining again a pair of acrocentric autosomes, painted majority of neo-Y, leaving a narrow centromere-proximal region on the long arms unstained. A probable X_1_ sex chromosome had only the proximal one third of its lenght marked by the NGU-Y probe, while the rest of the long arms was composed of DAPI-positive block. The NLO-Y probe painted entire neo-Y chromosome, including the narrow NGU-Y-negative gap. NLO-Y and NGU-Y probes also co-localized on the proximal half on X_1_. Additionally, the NLO-Y probe produced the signal patterns over many other chromosomes, some of which probably reflect rather repetitive content than identity of synteny blocks. Based on the neo-Y-linked patterns, one of the painted small chromosomes should be X_2_. Integrated WCP/BAC-FISH experiments confirmed the placement of *amhr2* in the (peri)centromeric region of neo-Y, and proximally on the long arms of X_1_. The THI-triv probe labelled one pair of small submeta-to-subtelocentric chromosomes, long arms of a pair of medium-sized subtelocentrics, and a narrow terminal portion of long arms on the pair of another medium-sized subtelocentrics.

Zoo-FISH in *N. ditte* produced exceptional patterns in the way that none of the applied probes matched the metacentric neo-Y. While X chromosomes could not be unambiguously identified, it may be inferred that the hybridization signals are absent also from them. The FTH- triv probe entirely painted one medium-sized submetacentric pair, and most of the length (except for a narrow terminal part on the long arms) of the medium-sized subtelocentric pair. The NGU-Y probe stained two submetacentric pairs, with NLO-Y probe co-hybridizing to the smaller from these pairs, but only across its long arms. Besides that, NLO-Y entirely painted one large, and one small acrocentric pair.

## Discussion

In this work, we studied four *Nothobranchius* killifish species and their outgroup *F. thierryi*, all having with X_1_X_2_Y sex chromosome system, using a combination of cytogenetic and genomic approaches. Given the high incidence of multiple sex chromosomes in the genus *Nothobranchius* (N=6), equaled among teleosts only by Neotropical *Harttia* catfishes (Deon et al. 2020; Sember et al. 2021) and well-resolved phylogenetic relationships (van der Merwe et al. 2021; Bartakova et al. in rev.), *Nothobranchius* killifishes are a highly informative lineage for investigating evolutionary forces that drive sex chromosome turnovers, particularly those occurring via sex chromosome-autosome fusions (Vicoso, 2019; Sember et al., 2021; Kitano et al. 2024). By applying sex chromosome-specific WCP probes from *N. guentheri*, *N. lourensi* and *F. thierryi*, and BAC-FISH with orthologues of usual fish MSD genes, we revealed at least four independent origins of sex chromosomes in this genus.

Specifically, X_1_X_2_Y systems of *N. guentheri*, *N. brieni* and *N. lourensi* share a synteny block but with independent autosomal additions. The shared synteny block of the X_1_X_2_Y system suggests possible ancestral sex chromosome pair, which originates from the common ancestor of these three species. This event would therefore precede the split between Coastal and Inland clades dated around 9-11 million years ago (Mya) (van der Merwe et al. 2021). It seems reasonable to assume that other closely related *Nothobranchius* species within these clades possess the same sex chromosomes which just cannot be revealed cytogenetically. To test homeology of the putative ancestral sex chromosome synteny block it will be critical to compare the patterns of sequence divergence and to calculate divergence times between sex chromosomes of these related species (cf. Charlesworth 2021; Li et al. 2021; Sardell et al. 2021).

We further show that formerly characterized XY sex chromosomes of *N. furzeri* and *N. kadleci* (Reichwald et al. 2015; Willemsen et al. 2020; Štundlová et al. 2022; Richter et al. 2023) share partial homology with the X_1_X_2_Y system in *F. thierryi*. The shared segment carries the *gdf6* gene which makes it a candidate MSD for *F. thierryi*. If confirmed, *gdf6* was likely co-opted independently from the system, which evolved in *N. furzeri* and *N. kadleci* after their divergence from *N. orthonotus* (Štundlová et al. 2022). Its independent co-option may point to phylogenetic constraint in sex determination of killifishes as *gdf6* has been thus far identified as putative MSD only in the Pachón cave population of blind tetra *Astyanax mexicanus* (Imarazene et al. 2021) and it is among potential candidates for MSD in Greenland Halibut (Ferchaud et al. 2022).

We also demonstrate that X_1_X_2_Y sex chromosomes of *N. ditte* evolved independently from yet another linkage groups, which are not involved in sex chromosomes of any other studied *Nothobranchius* species. The sex in this species is therefore likely determined by yet another MSD. As we tested most of the usual suspects for the MSD role in teleosts by means of BAC FISH (Herpin & Schartl 2015; Pan et al. 2021; Kitano et al. 2024), sex in *N. ditte* can be determined either by gene duplicate or novel mechanisms.

Reverse-FISH WCP experiments in all studied species confirmed previous assumptions (Krysanov et al. 2016; Krysanov & Demidova 2018) that the *Nothobranchius* multiple sex chromosome systems evolved by Y-autosome fusion. WCP probes painted equally all the members of the X_1_X_2_Y system from end to end, which may imply low molecular differentiation between gametologs. The same WCP pattern was, however, found also in *Oplegnathus* knifefishes (Xu et al. 2019), where subsequent genomic study showed moderate neo-Y degeneration (Li et al. 2021).

The cross-species WCP patterns further imply complex evolutionary history of neo-sex chromosomes in *N. lourensi* and *N. brieni*. While the large subtelocentric neo-Y chromosome of *N. lourensi* originated from subsequent fusions of three different synteny blocks, the neo-Y of *N. brieni* evolved through fusion and subsequent inversion, which was indicated by disjoint arrangement of segments painted by the NGU-Y probe (an opposite order of events than documented e.g. in X_1_X_2_Y multiple sex chromosome system of *Oplegnathus punctatus*; Li et al. 2021). Furthermore, the comparison of the hybridization patterns of *amhr2* (BAC-FISH) and rDNA clusters on X_1_ and neo-Y chromosome of *N. guentheri* (Supplementary Figs 1 and 2) suggested subsequent small-scale inversion(s) following the original fusion, which shifted the order of these regions and the position of centromere on the neo-Y. It is worth mentioning that NGU-Y probe confirmed the identity and composition of multiple sex chromosomes in both studied *N. guentheri* populations, which, however, differ by their 18S rDNA FISH patterns.

To determine degree of sex chromosome differentiation extent of non-recombining regions, and potential MSD genes, we performed long-read sequencing of the *F. thierryi* and *N. guentheri* genomes, combined with coverage analysis and male vs. female Pool-seq data. As for the difference in coverage between male and female genomes, in both species, the sequences have not yet diverged enough, pointing to low to moderate level of X-Y differentiation in the non-recombining region.

In case of *F. thierryi*, the *gdf6* gene localizes within the non-recombining region. In *N. guentheri*, our analysis showed two clearly defined regions enriched for male-specific single nucleotide polymorphisms (SNPs), which differed in their degree of sequence divergence between gametologs. This is suggestive of the presence of two evolutionary strata i.e. regions which stopped recombining at different evolutionary time points (e.g. Lahn & Page 1999; Zhou et al. 2014). Evolutionary strata have been already found also among teleosts, e.g. in guppies (reviewed in Darolti et al. 2022) and sticklebacks (Peichel et al. 2020; Sardell et al. 2021). The older stratum may correspond to the ancestral sex-linked chromosome shared by the Coastal and Inland clades. The *amhr2* gene is, however, localized in the younger of the two strata and high similarity of its X and Y-linked alleles questions its possible role in sex determination in these species. Genes annotated in the older stratum do not provide any other clear candidate for MSD in *N. guentheri*. It is, nonetheless, of note, that a single SNP that differentiated Y-linked *amhr2* gene copy is located in the first exon which is consistent with the emerging role of mutations in its N-terminal domain crucial for ligand binding specifity in sex determination (Wen et al. 2022; Kuhl et al. 2023 and references therein).

Interestingly, male-determining regions in *F. thierryi*, *N. brieni, N. guentheri* and *N. lourensi* tend to locate close to centromere or close to an assumed rearrangement breakpoints on neo-Y, as cytologically confirmed by WCP and BAC-FISH with *gdf6* and *amhr2* orthologs (schematized in Fig. 3). An increasing number of studies evidenced significantly reduced recombination around the chromosome fusion junctions in various taxa (Gil-Fernández et al. 2020; Vara et al. 2021; Yoshida et al. 2023; MacLeod-Bigley & Boulding 2023; Lisachov et al. 2023; Renner et al. 2023), and several lines of cytogenetic or genomic evidence suggest this process might have contributed to sequence differentiation of multiple sex chromosomes of certain fish species (de Moraes et al. 2019; Li et al. 2021; Sardell et al. 2021). The effect of suppressed recombination might be further enhanced by the close proximity to centromere as it is widely known that pericentromeric regions display low recombination rates (Nambiar & Smith 2016; Miller & Hawley 2017; Filatov 2024). Last but not least, Haenel et al. (2018) found out that in species with large chromosomes, recombination rate is low in the central chromosome regions. All these mechanisms might have separately or in combination contributed to the observed patterns of differentiation of *Nothobranchius* sex-determining Y-linked regions. Particularly in *N. guentheri*, Y-autosome fusion might have an effect on creation on younger stratum, while the older stratum, which we consider as ancestral sex-linked region, should have formed before the fusion. The presence of sex-determining regions in the interstitial position on the large Y chromosomes may substantially contribute to reduced or abolished recombination in species with heterochiasmy (i.e. sex-specific differences in recombination landscapes; Sardell & Kirkpatrick 2020), which was proposed to occur in *N. furzeri* and *N. kadleci* with recombination restricted towards the chromosome ends in males (Štundlová et al. 2022). Hence, the proximity to centromere and/or heterochiasmy may be considered as important driving factors in the ancestral sex chromosome region (i. E. old stratum in *N. guentheri*) establishment. Further studies need to be performed to assess whether different X_1_X_2_Y systems in *Nothobranchius* also evolved under the effect of heterochiasmy.

Several hypotheses on mechanisms and forces driving sex chromosome turnover and its fixation have been put forward (Saunders 2019; Vicoso 2019) and they may be applied also on the specific situation of sex chromosome-autosome fusions (Pennell et al. 2015; Kitano et al. 2024). One possible trigger may be selection on the newly created linkage disequillibrium between sex chromosome and autosomal addition, which might resolve sexual conflict (Charlesworth & Charlesworth 1980; van Doorn & Kirkpatrick 2007; Kitano et al. 2009; Matsumoto & Kitano 2016). Other models rely on action of genetic drift (Bull & Charnov 1977; Veller et al. 2017, Saunders et al. 2018) and the load of deleterious mutations (Blaser et al. 2013, 2014). In *Nothobranchius*, the turnover rate may have been facilitated by their often small effective population sizes (Bartáková et al. 2013; Cui et al. 2019; van der Merwe et al. 2021). This is consistent with the action of both genetic drift and mutation load model, (Blaser et al. 2014; Saunders et al. 2018) and also with the maintenance of male heterogamety (Jeffries et al. 2018; Saunders 2019).

Last but not least, our study highlights the complementarity between cytogenetic and genomic approaches in studying sex chromosome evolution. In contrast to some highly degenerated systems e.g. in insects (Vicoso & Bachtrog 2015), fish multiple sex chromosomes show mostly little differentiation (Sember et al. 2021). Yet, there have been few fish multiple sex chromosome systems traced both by cytogenetic and genomic methods, as exemplified by *Gasterosteus* sticklebacks (e.g. Kitano et al. 2009; Sardell et al. 2021; Dagilis et al. 2022), *Oplegnathus* knifefishes (Xu et al. 2019; Xiao et al. 2020; Li et al. 2021; Gong et al. 2022), and a spinyhead croker *Collichthys lucidus* (Zhang et al. 2018; Cai et al. 2019).

## Conclusions

*Nothobranchius* sex chromosomes are highly dynamic and evolved repeatedly and recurrently from different linkage groups. Taking into account the availability of well-resolved phylogeny, this fish genus represents excellent model system to investigate fundamental questions related to sex chromosome evolution and turnover. Our findings in the present study collectively suggest at least four independent origins of sex chromosomes – one in outgroup *F. thierryi* and three in the genus *Nothobranchius*: i) in the common ancestor of *N. furzeri* and *N. kadleci*, ii) before the split of Coastal and Inland clade and iii) in *N. ditte*. We also delineated regions of recombination suppression in *F. thierryi* and *N. guentheri* and specifically in the latter we found evidence for two evolutionary strata. We suppose a possible importance of centromeric regions and fusion points in facilitating recombination suppression for development and spreading of sex-determining region. Our study highlights the need for combined cytogenetic and genomic methodologies in analysing sex chromosomes. These data are mutually informative and allowed to draw a more complete picture of *Nothobranchius* sex chromosome evolution.

## Supporting information

Hospodarska_et_al_2024_Supplementary_Table_2

## Acknowledgements

We gratefully acknowledge B. Nagy and H. Hengstler for providing part of the study material and A. Nikiforov for his help in breeding and keeping fishes. We are also grateful to P. Šejnohová for her laboratory assistance.

## Author Contributions

Conceptualization: AS, PN; Data curation: PN; Formal analysis: MHo, PM, ACV, PN; Funding acquisition: AS, CE; Investigation: MHo, PM, ACV, AA-R, SAS, TP, MA, KJ, JŠ, MJ, MHi, TL, EYuK, PN, AS; Methodology: AS, PN, ACV, AA-R, TL, SAS; Project administration: AS, PN; Resources: AS, PN, MR, CE, PR; Supervision: AS, PN, MR; Validation: MHo, PM, ACV, MA, AS, PN; Writing original draft: AS, MHo, PN, PM; Writing—review & editing: PN, AS, PM, MA, MR, ACV, MHo, CE, PR, SAS, EYuK.

## Statement of Ethics

The experimental part involving fish individuals was supervised by the Institutional Animal Care and Use Committee of the Institute of Animal Physiology and Genetics CAS, v.v.i., with the supervisor’s permit number CZ 02361 certified and issued by the Ministry of Agriculture of the Czech Republic. The handling of fish individuals to obtain chromosomes followed European standards in agreement with §17 of the Act No. 246/1992 coll. The experiments with *N. brieni*, *N. ditte*, and one population of *N. guentheri* (NGU2) were approved by the Ethics Committee of Severtsov Institute of Ecology and Evolution (Order No. 27 of November 9, 2018). In case of chromosome preparations from the kidney and gonads, fishes were euthanized before organ sampling using 2-phenoxyethanol (Sigma-Aldrich, St. Louis, MO, USA). A narrow stripe of the tail fin was taken from live specimens after fishes were anaesthetized by MS-222 (Merck KGaA, Darmstadt, Germany).

## Supplementary material

**Supplementary Table 1.**
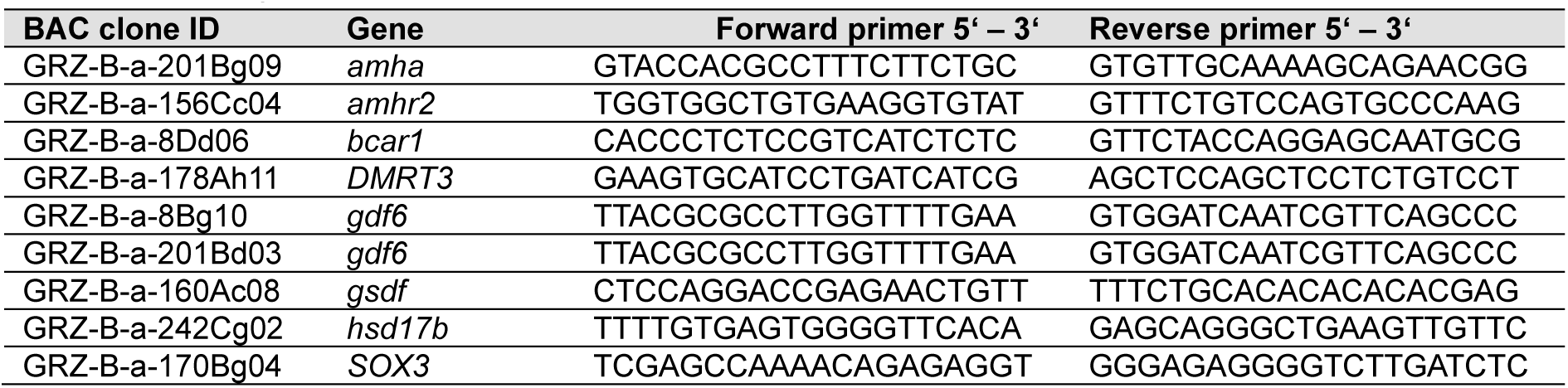
Primers for BAC clone verification.

**Supplementary Table 2**. *Fundulosoma thierryi* and *Nothobranchius guentheri* long-read genome sequencing, raw data – disclosed as separate file.

**Supplementary Fig. 1.**
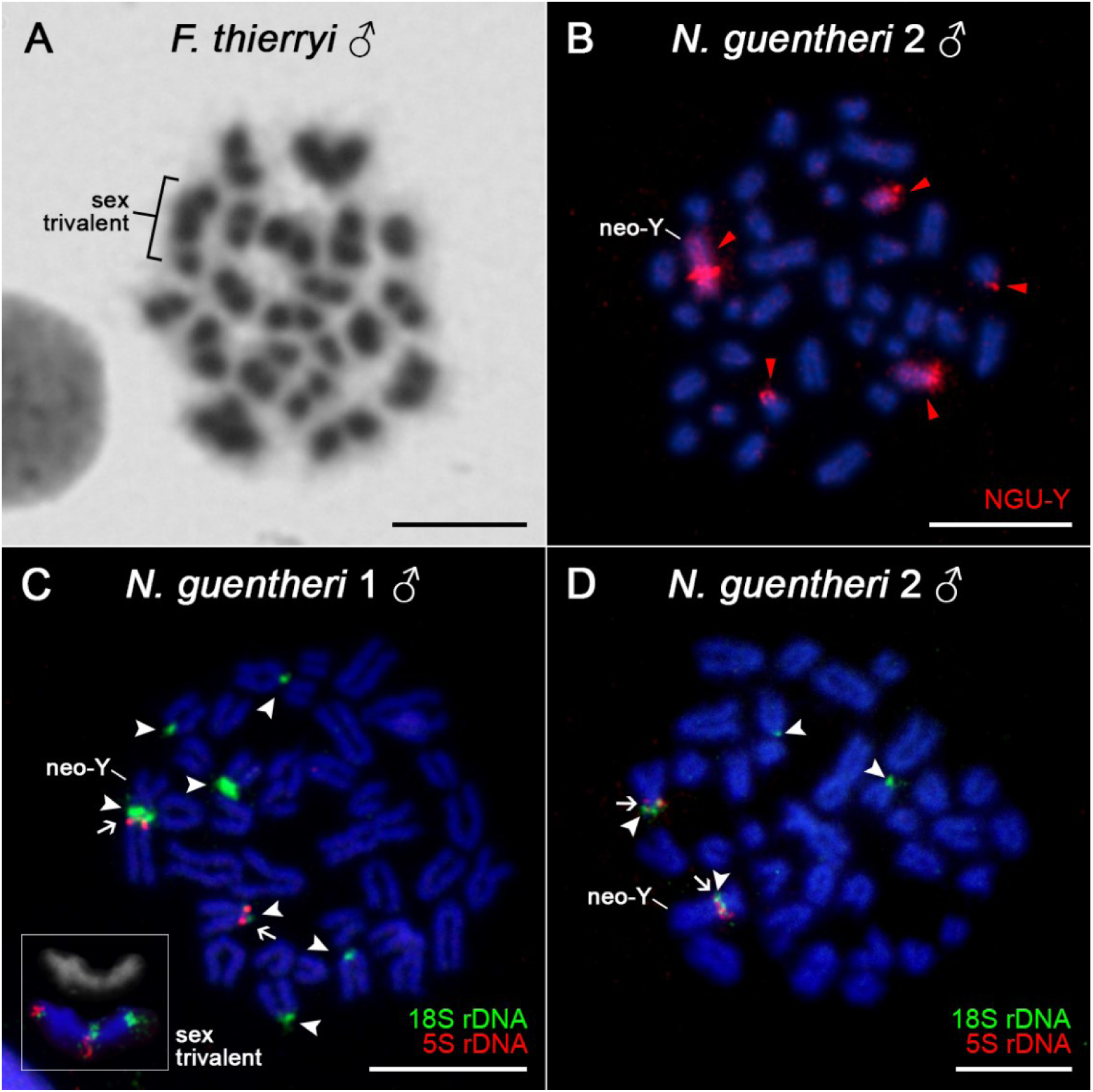
A collection of additional supportive cytogenetic results. **A**) An example of meiotic spread of *F. thierryi* male used for microdissection procedure. A detectable sex-trivalent which has been cut out for the painting probe construction is depicted. **B**) WCP with NGU-Y probe generated from NGU1 population on the male mitotic metaphase of NGU2 individual, confirming the identity of all three sex chromosomes also in this population. Note additional signals on the short arms of two acrocentic chromosomes. **C-D**) rDNA FISH hybridization patterns in *N. guentheri* males of population NGU1 and NGU2, respectively. Dual-colour FISH with 5S (red signals; arrows) and 18S (green signals; arrowheads) rDNA probes. Chromosomes were counterstained with DAPI (blue). Note that 18S rDNA clusters occupy short arms of six acrocentric or subtelocentric chromosomes (including X chromosomes), and interstitial region on long arms of the neo-Y chromosome in NGU1 population, while it is present on four chromosomes in NGU2 population, including neo-Y and one X chromosome. The signals on two acrocentric autosomes highly probably correspond with WCP signals in **B**). In the population NGU1 (**C**), we probed also meiotic plates and a trivalent formed from X_1_X_2_Y sex chromosomes is herein framed, along with rDNA signal patterns confirming that all three members of the system carry one or both rDNA clusters. 5S rDNA probe showed, in contrast to the 18S rDNA one, the same patterns in both populations, as it overlaps with the 18S rDNA cluster on neo-Y chromosome and one X chromosome. Scale bar = 10 μm.

**Supplementary Fig. 2.**
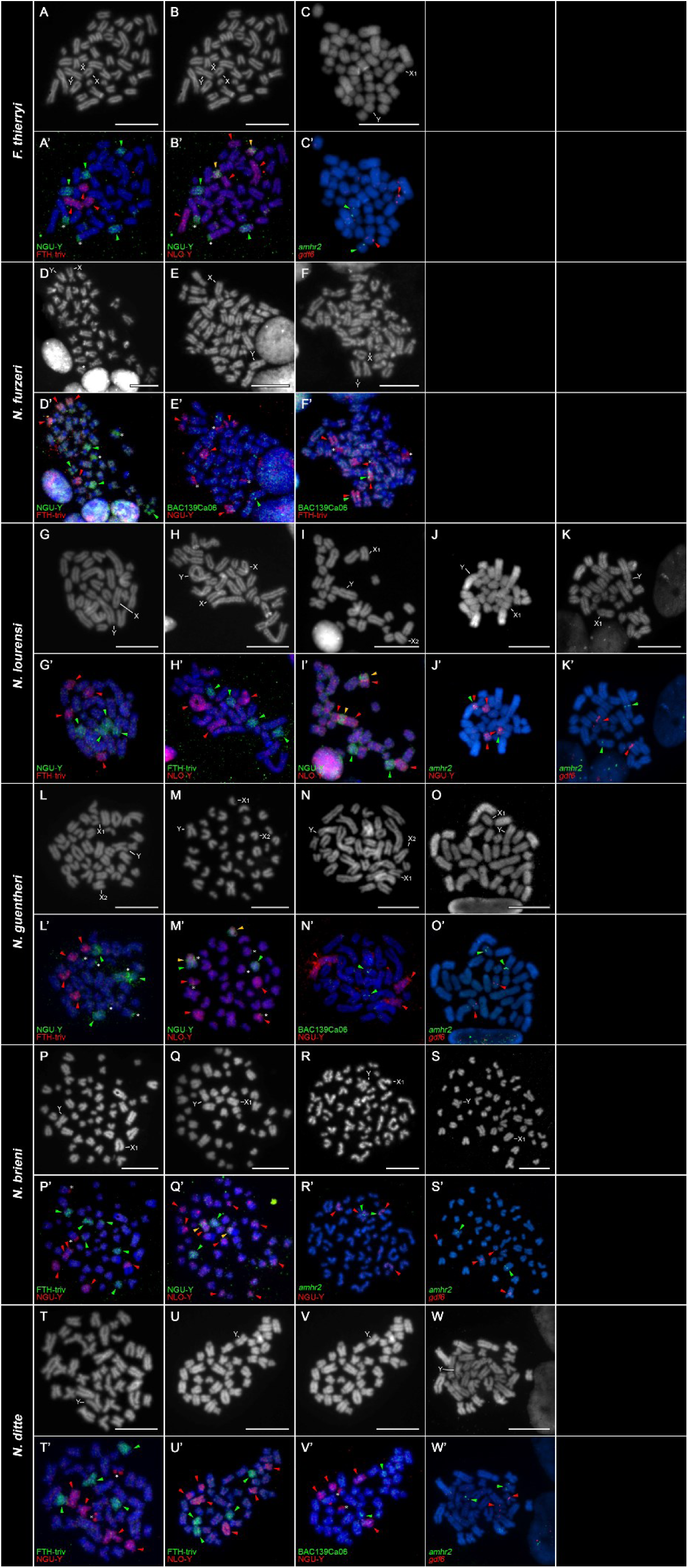
Male mitotic metaphases of studied *Nothobranchius* and *Fundulosoma* species after cross-species experiments with WCP and BAC-FISH probes. To improve clarity of the signal patterns, some included metaphases were reprobed for two different experiments. For each species, upper row represents DAPI images, converted to grayscale, while bottom row shows merged FISH pictures with two mapped fluorescent probes (green and red signals; arrowheads) and DAPI (blue). Sex chromosome-derived probes: NGU-Y (*N. guentheri*; Y chromosome), NLO-Y (*N. lourensi*; Y chromosome), FTH-triv (*F. thierryi*, whole meiotic sex-trivalent X_1_X_2_Y). BAC-FISH: clones derived from *N. furzeri* BAC library carrying *gdf6* and *amhr2* gene, and Y-linked 139Ca06 clone (**Reichwald et al. 2015**). Sex chromosomes are marked if possible (see upper DAPI images). Due to sterical reasons, neo-Y is marked as “Y”. X chromosomes are distinguished as X_1_ (ancestral) or X_2_ (new autosome addition; neo-X) if the signal patterns allow that. Asterisks denote additional signals which, based on the complementary data, represent major ribosomal DNA (rDNA) sites whose tandem arrays are located also on the neo-Y of *N. guentheri* (hence being included in the painting probe). Note that painting probes fully stain all members of the X_1_X_2_Y system from which they were derived (in *F. thierryi*, *N. guentheri*, *N lourensi*). In all analyzed species, NGU-Y and NLO-Y probes partially overlap (orange arrowheads) in one segment (*N. guentheri*, *N. lourensi*, *N. brieni*) or by covering entire autosome pair (*F. thierryi*), or its short arms (*N. ditte*). Neo-Y linked regions stained both by NGU-Y and NLO-Y probes also carry *amhr2* BAC-FISH signal. Notice also that *F. thierryi* and *N. furzeri* share the synteny block carrying *gdf6* gene (on the long arms of the Y chromosome in *N. furzeri*, and close to centromere on neo-Y of *N. lourensi*). In *N. ditte*, none of the mapped probes produced signals on neo-Y. Scale bar = 10 μm.

## References

Abbott JK, Nordén AK, Hansson B. (2017). Sex chromosome evolution: historical insights and future perspectives. Proc R Soc B Biol Sci 284: 20162806. 10.1098/rspb.2016.2806

Al-Rikabi A, Liehr LB, Liehr T. (2020). Glass-needle based chromosome microdissection – how to set up probes for molecular cytogenetics? Video Journal of Clinical Research 2:100004VAM08AR2020. 10.5348/100004VAM08AR2020TR

Alonge M, Soyk S, Ramakrishnan S, Wang X, Goodwin S, Sedlazeck FJ, Lippman ZB, Schatz MC. (2019). RaGOO: fast and accurate reference-guided scaffolding of draft genomes. Genome Biology 20: 224. 10.1186/s13059-019-1829-6

Andrews S (2010). FastQC: a quality control tool for high throughput sequence data. http://www.bioinformatics.babraham.ac.uk/projects/fastqc/. Accessed 25 Sept 2019

Bartakova V, Bryjová A, Polačik M, Alila D, Nagy B, Watters B, Bellstedt D, Blažek R, Žák J, Reichard M. Phylogenomics of *Nothobranchius* in lowland Tanzania: species delimitation and comparative genetic structure. Molecular Phylogenetics and Evolution in revision

Bartáková V, Reichard M, Janko K, Polačik M, Blažek R, Reichwald K, Cellerino A, Bryja J. (2013). Strong population genetic structuring in an annual fish, *Nothobranchius furzeri*, suggests multiple savannah refugia in southern Mozambique. BMC Evolutionary Biology 13: 196.

Bartáková V, Reichard M, Blažek R, Polačik M, Bryja J. (2015). Terrestrial fishes: Rivers are barriers to gene flow in annual fishes from the African savanna. Journal of Biogeography 42: 1832–1844. 10.1111/jbi.12567

Bertollo LAC, Cioffi MB, Moreira-Filho O. (2015). Direct chromosome preparation from freshwater teleost fishes. In: Ozouf-Costaz C, Pisano E, Foresti F, and de AlmeidaToledo LF (eds) Fish cytogenetic techniques ray-fin fishes and chondrichthyans. CRC Press, Inc, Endfield, pp 21–26. 10.1201/b18534-4

Blaser O, Grossen C, Neuenschwander S, Perrin N. (2013). Sex-chromosome turnovers induced by deleterious mutation load. Evolution 67: 635–645. 10.1111/j.1558-5646.2012.01810.x

Blaser O, Neuenschwander S, Perrin N. (2014). Sex-chromosome turnovers: the hot-potato model. The American Naturalist 183:140–146. 10.1086/674026

Blažek R, Polačik M, Kačer P, Cellerino A, Řežucha R, Methling C, Tomášek O, Syslová K, Terzibasi Tozzini E, Albrecht T, Reichard M. (2017). Repeated intraspecific divergence in life span and aging of African annual fishes along an aridity gradient. Evolution 71: 386–402. 10.1111/evo.13127

Bolger AM, Lohse M, Usadel B. (2014). Trimmomatic: A flexible trimmer for Illumina sequence data. Bioinformatics 30: 2114–2120. 10.1093/bioinformatics/btu170

Brůna T, Hoff KJ, Lomsadze A, Stanke M, Borodovsky M. (2021). BRAKER2: Automatic eukaryotic genome annotation with GeneMark-EP+ and AUGUSTUS supported by a protein database. NAR Genomics and Bioinformatics 3: lqaa108. 10.1093/nargab/lqaa108

Bull JJ, Charnov EL. (1977). Changes in the heterogametic mechanism of sex determination. Heredity 39: 1–14. 10.1038/hdy.1977.38

Cellerino A, Valenzano DR, Reichard M. (2016). From the bush to the bench: The annual *Nothobranchius* fishes as a new model system in biology. Biological Reviews of the Cambridge Philosophical Society 91: 511–533. 10.1111/brv.12183

Chan PP, Lowe TM. (2019). tRNAscan-SE: searching for tRNA genes in genomic sequences. In M. Kollmar (Ed.), Gene prediction. Methods in molecular biology, vol. 1962 (pp. 1– 14). Humana. 10.1007/978-1-4939-9173-0_1

Charlesworth D. (2021). The timing of genetic degeneration of sex chromosomes. Philosophical Transactions of the Royal Society B 376: 20200093. 10.1098/rstb.2020.0093

Charlesworth D, Charlesworth B. (1980). Sex differences in fitness and selection for centric fusions between sex-chromosomes and autosomes. Genetics Research 35: 205–214. 10.1017/S0016672300014051

Charlesworth D, Charlesworth B, Marais G. (2005). Steps in the evolution of heteromorphic sex chromosomes. Heredity 95:118–128. 10.1038/sj.hdy.6800697

Cioffi MB, Yano CF, Sember A, Bertollo, LAC. (2017). Chromosomal evolution in lower vertebrates: sex chromosomes in Neotropical fishes. Genes 8: 258. 10.3390/genes8100258

Cortez D, Marin R, Toledo-Flores D, Froidevaux L, Liechti A, Waters PD, Grützner F, Kaessmann H. (2014). Origins and functional evolution of Y chromosomes across mammals. Nature 508: 488–493. 10.1038/nature13151

Cui R, Medeiros T, Willemsen D, Iasi LNM, Collier GE, Graef M, Reichard M, Valenzano DR. (2019). Relaxed selection limits lifespan by increasing mutation load. Cell 178: 385– 399.e20. 10.1016/j.cell.2019.06.004

Dagilis AJ, Sardell JM, Josephson MP, Su Y, Kirkpatrick M, Peichel CL. (2022). Searching for signatures of sexually antagonistic selection on stickleback sex chromosomes. Philosophical Transactions of the Royal Society B 377: 20210205.

Darolti I, Almeida P, Wright AE, Mank JE. (2022). Comparison of methodological approaches to the study of young sex chromosomes: A case study in *Poecilia*. Journal of Evolutionary Biology, 35: 1646–1658. 10.1111/jeb.14013

De Coster W, D’Hert S, Schultz DT, Cruts M, Van Broeckhoven C. (2018). NanoPack: Visualizing and processing long-read sequencing data. Bioinformatics 34: 2666–2669. 10.1093/bioinformatics/bty149

de Moraes RL, Sember A, Bertollo LA, de Oliveira EA, Ráb P, Hatanaka T, Marinho MMF, Liehr T, Al-Rikabi ABH, Feldberg E, Viana PF, Cioffi MB. (2019). Comparative cytogenetics and neo-Y formation in small-sized fish species of the genus *Pyrrhulina* (Characiformes, Lebiasinidae). Frontiers in Genetics 10: 678. 10.3389/fgene.2019.00678

Deon GA, Glugoski L, Vicari MR, Nogaroto V, Sassi FMC, Cioffi MB, Liehr T, Bertollo LAC, Moreira-Filho O. (2020). Highly rearranged karyotypes and multiple sex chromosome systems in armored catfishes from the genus *Harttia* (Teleostei, siluriformes). Genes 11: 1366. 10.3390/genes11111366

Dobin A, Gingeras TR. (2015). Mapping RNA-seq reads with STAR. Current Protocols in Bioinformatics 51: 11–14. 10.1002/0471250953.bi1114s51

Dorn A, Musilová Z, Platzer M, Reichwald K, Cellerino A. (2014). The strange case of East African annual fish: Aridification correlates with diversification for a savannah aquatic group? BMC Evolutionary Biology 14: 210. 10.1186/s12862-014-0210-3

El Taher A, Ronco F, Matschiner M, Salzburger W, Böhne A. (2021). Dynamics of sex chromosome evolution in a rapid radiation of cichlid fishes. Science Advances 7: eabe8215. 10.1126/sciadv.abe8215

Ewulonu UV, Haas R, Turner BJ. (1985). A multiple sex chromosome system in the annual killifish, *Nothobranchius guentheri*. Copeia 1985: 503–508. 10.2307/1444868

Ferchaud A-L, Mérot C, Normandeau E, Ragoussis J, Babin C, Djambazian H, Bérubé P, Audet C, Treble M, Walkusz W, Bernatchez L. (2022). Chromosome-level assembly reveals a putative Y-autosomal fusion in the sex determination system of the Greenland Halibut (*Reinhardtius hippoglossoides*). G3 (Bethesda) 12: jkab376. 10.1093/G3JOURNAL/JKAB376

Feron R, Jaron KS. (2021). SexGenomicsToolkit/PSASS: 3.1.0. 10.5281/zenodo.4442702

Filatov DA. (2024). Evolution of a plant sex chromosome driven by expanding pericentromeric recombination suppression. Scientific Reports 14: 1373. 10.1038/s41598-024-51153-0

Flynn JM, Hubley R, Goubert C, Rosen J, Clark AG, Feschotte C, Smit AF. (2020). RepeatModeler2 for automated genomic discovery of transposable element families. Proceedings of National Academy of Sciences USA 117: 9451–9457. 10.1073/pnas.1921046117

Fricke R, Eschmeyer WN, & Van der Laan R (eds) (2024). Eschmeyer’s catalog of fishes: genera, species, references. http://researcharchive.calacademy.org/research/ichthyology/catalog/fishcatmain.asp. Accessed 3 March 2024

Furness AI. (2016). The evolution of an annual life cycle in killifish: Adaptation to ephemeral aquatic environments through embryonic diapause. Biological Reviews of the Cambridge Philosophical Society 91: 796–812. 10.1111/brv.12194

Gamble T. (2016). Using RAD-seq to recognize sex-specific markers and sex chromosome systems. Molecular Ecology 25: 2114–2116. 10.1111/mec.13648

Gil-Fernández A, Saunders PA, Martín-Ruiz M, Ribagorda M, López-Jiménez P, Jeffries DL, et al. (2020). Meiosis reveals the early steps in the evolution of a neo-XY sex chromosome pair in the African pygmy mouse *Mus minutoides*. PLoS Genetics 16: e1008959. 10.1371/journal.pgen.1008959

Gong J, Li B, Zhao J, Ke Q, Xu D, Zhou T, Xu P. (2022). Sex-specific genomic region identification and molecular sex marker development of rock bream (*Oplegnathus fasciatus*). Marine Biotechnology 24:163–173. 10.1007/s10126-022-10095-2

Guan D, McCarthy SA, Wood J, Howe K, Wang Y, Durbin R. (2020). Identifying and removing haplotypic duplication in primary genome assemblies. Bioinformatics 36: 2896–2898. 10.1093/bioinformatics/btaa025

Guiguen Y, Fostier A, Herpin A. (2018). Sex determination and differentiation in fish: genetic, genomic, and endocrine aspects. In: Wang HP, Piferrer F, Chen S-L (eds) Sex Control in Aquaculture, 1st edn. John Wiley & Sons, Hoboken, pp 35–63

Gurevich A, Saveliev V, Vyahhi N, Tesler G. (2013). QUAST: quality assessment tool for genome assemblies. Bioinformatics 29: 1072–1075. 10.1093/bioinformatics/btt086

Haenel Q, Laurentino TG, Roesti M, Berner D. (2018). Meta-analysis of chromosome-scale crossover rate variation in eukaryotes and its significance to evolutionary genomics. Molecular Ecology 27: 2477–2497. 10.1111/mec.14699

Herpin A, Schartl M. (2015). Plasticity of gene-regulatory networks controlling sex determination: of masters, slaves, usual suspects, newcomers, and usurpators. EMBO Rep. 16: 1260–1274. 10.15252/embr.201540667

Holley G, Beyter D, Ingimundardottir H, Møller PL, Kristmundsdottir S, Eggertsson HP, Halldorsson BV. (2021). Ratatosk: Hybrid error correction of long reads enables accurate variant calling and assembly. Genome Biology 22: 28. 10.1186/s13059-020-02244-4

Hu CK, Brunet A. (2018). The African turquoise killifish: A research organism to study vertebrate aging and diapause. Aging Cell 17: e12757. 10.1111/acel.12757

Hu J, Fan J, Sun Z, Liu S. (2020). NextPolish: A fast and efficient genome polishing tool for long-read assembly. Bioinformatics 36: 2253–2255. 10.1093/bioinformatics/btz891

Humann JL, Lee T, Ficklin S, Main D. (2019). Structural and functional annotation of eukaryotic genomes with GenSAS. In M. Kollmar (Ed.), Gene Prediction. Methods in Molecular Biology, vol. 1962. Humana. 10.1007/978-1-4939-9173-0_3

Imarazene B, Du K, Beille S, Jouanno E, Feron R, Pan Q, Torres-Paz J, Lopez-Roques C, Castinel A, Gil L, et al. (2021). A supernumerary “B-sex” chromosome drives male sex determination in the Pachón cavefish, *Astyanax mexicanus*. Current Biology 31: 4800– 4809.e9. 10.1016/j.cub.2021.08.030

Jeffries DL, Lavanchy G, Sermier R, Sredl MJ, Miura I, Borzée A, Barrow LN, Canestrelli D, Crochet P-A, Dufresnes C, et al. (2018) A rapid rate of sex-chromosome turnover and non-random transitions in true frogs. Nature Communications 9: 4088. 10.1038/s41467-018-06517-2

Kitano J, Ross JA, Mori S, Kume M, Jones FC, Chan YF, Absher DM, Grimwood J, Schmutz J, Myers RM, Kingsley DM, Peichel CL. (2009). A role for a neo-sex chromosome in stickleback speciation. Nature 461:1079–1083. 10.1038/nature08441

Kitano J, Peichel CL. (2012). Turnover of sex chromosomes and speciation in fishes. Environmental Biology of Fishes 94: 549–558. 10.1007/s10641-011-9853-8

Kitano J, Ansai S, Takehana Y, Yamamoto Y. (2024). Diversity and convergence of sex determination mechanisms in teleost fish. Annual Review of Animal Biosciences 12: 233–259. 10.1146/annurev-animal-021122-113935

Kligerman AD, Bloom SE. (1977). Rapid chromosome preparations from solid tissues of fishes. Journal of the Fish Research Board of Canada 34: 266–269. 10.1139/f77-039

Kolmogorov M, Yuan J, Lin Y, Pevzner PA. (2019). Assembly of long, error-prone reads using repeat graphs. Nature Biotechnology 37: 540–546. 10.1038/s41587-019-0072-8

Kratochvíl L, Stöck M, Rovatsos M, Bullejos M, Herpin A, Jeffries DL, Peichel CL, Perrin N, Valenzuela N, Johnson Pokorná M. (2021). Expanding the classical paradigm: what we have learnt from vertebrates about sex chromosome evolution. Philosophical Transactions of the Royal Society B 376: 20200097. 10.1098/rstb.2020.0097

Krysanov E, Demidova T, Nagy B. (2016). Divergent karyotypes of the annual killifish genus *Nothobranchius* (Cyprinodontiformes, Nothobranchiidae). Comparative Cytogenetics 10: 439–445. 10.3897/CompCytogen.v10i3.9863

Krysanov E, Demidova T. (2018). Extensive karyotype variability of African fish genus *Nothobranchius* (Cyprinodontiformes). Comparative Cytogenetetics 12: 387–402. 10.3897/CompCytogen.v12i3.25092

Krysanov EY, Nagy B, Watters BR, Sember A, Simanovsky SA. (2023). Karyotype differentiation in the *Nothobranchius ugandensis* species group (Teleostei, Cyprinodontiformes), seasonal fishes from the east African inland plateau, in the context of phylogeny and biogeography. Comparative Cytogenetics 7: 13–29. 10.3897/compcytogen.v7.i1.97165

Kuhl H, Euclide PT, Klopp C, Cabau C, Zahm M, Rogues C, Iampietro C, Kuchly C, Donnadieu C, Feron R, et al. (2023). Multi-genome comparisons reveal gain-and-loss evolution of the anti-Mullerian hormone receptor type 2 gene, an old master sex determining gene, in Percidae. Preprint (bioRxiv) 10.1101/2023.11.13.566804)

Kuhl H, Guiguen Y, Höhne C, Kreuz E, Du K, Klopp C, Lopez-Roques C, Yebra Pimentel ES, Ciorpac M, Gessner J, et al. (2021). A 180 Myr-old female-specific genome region in sturgeon reveals the oldest known vertebrate sex determining system with undifferentiated sex chromosomes. Philosophical Transactions of the Royal Society B 376: 20200089.

Lahn BT, Page DC. (1999). Four evolutionary strata on the human X chromosome. Science 286: 964–967. 10.1126/science.286.5441.964

Laetsch DR, Blaxter ML. (2017). BlobTools: Interrogation of genome assemblies. F1000Research 6: 1287. 10.12688/f1000research.12232.1

Lagesen K, Hallin P, Rødland EA, Stærfeldt H-H, Rognes T, Ussery DW. (2007). RNAmmer: Consistent and rapid annotation of ribosomal RNA genes. Nucleic Acids Research 35: 3100–3108. 10.1093/nar/gkm160

Levan AK, Fredga K, Sandberg AA. (1964). Nomenclature for centromeric position on chromosomes. Hereditas 52: 201–220. 10.1111/j.1601-5223.1964.tb01953.x

Li H, Durbin R. (2009). Fast and accurate short read alignment with Burrows–Wheeler transform. Bioinformatics 25: 1754–1760. 10.1093/bioinformatics/btp324

Li H, Handsaker B, Wysoker A, Fennell T, Ruan J, Homer N, Marth G, Abecasis G, Durbin R. (2009). The sequence alignment/map format and SAMtools. Bioinformatics 25: 2078– 2079. 10.1093/bioinformatics/btp352

Li M, Zhang R, Fan G, Xu W, Zhou Q, Wang L, Li W, Pang Z, Yu M, Liu Q, Liu X, Schartl M, Chen S. (2021). Reconstruction of the origin of a neo-Y sex chromosome and its evolution in the spotted knifejaw, *Oplegnathus punctatus*. Molecular Biology and Evolution 38: 2615–2626. 10.1093/molbev/msab056

Lisachov A, Tishakova K, Romanenko S, Lisachova L, Davletshina G, Prokopov D, Kratochvíl L, O’Brien P, Ferguson-Smith M, Borodin P, Trifonov V. (2023). Robertsonian fusion triggers recombination suppression on sex chromosomes in *Coleonyx* geckos. Scientific Reports 13: 15502. 10.1038/s41598-023-39937-2

Lomsadze A, Burns PD, Borodovsky M. (2014). Integration of mapped RNA-Seq reads into automatic training of eukaryotic gene finding algorithm. Nucleic Acids Research 42: e119. 10.1093/nar/gku557

Lukšíková K, Pavlica T, Altmanová M, Štundlová J, Pelikánová Š, Simanovsky SA, Yu KE, Jankásek M, Hiřman M, Reichard M, Ráb P, Sember A. (2023). Conserved satellite DNA motif and lack of interstitial telomeric sites in highly rearranged African Nothobranchius killifish karyotypes. Journal of Fish Biology 103: 1501–1514. 10.1111/jfb.15550

MacLeod-Bigley MLM, Boulding EG. (2023). High-density linkage maps detail sex-specific regions of suppressed recombination near fusions of polymorphic chromosomes in purebred and hybrid North American Atlantic salmon (*Salmo salar* L.). Genome 66: 175–192. 10.1139/gen-2022-0065

Manni M, Berkeley MR, Seppey M, Simão FA, Zdobnov EM. (2021). BUSCO update: Novel and streamlined workflows along with broader and deeper phylogenetic coverage for scoring of eukaryotic, prokaryotic, and viral genomes. Molecular Biology and Evolution, 38: 4647–4654. 10.1093/molbev/msab199

Marçais G, Kingsford C. (2011). A fast, lock-free approach for efficient parallel counting of occurrences of k-mers. Bioinformatics 27 : 764–770. 10.1093/bioinformatics/btr011

Martin M. (2011). Cutadapt removes adapter sequences from high-throughput sequencing reads. EMBnet J 17: 10–12. 10.14806/ej.17.1.200

Matsumoto T, Kitano J. (2016). The intricate relationship between sexually antagonistic selection and the evolution of sex chromosome fusions. Journal of Theoretical Biology 404: 97–108. 10.1016/j.jtbi.2016.05.036

Meisel RP. (2020). Evolution of sex determination and sex chromosomes: a novel alternative paradigm. BioEssays 42: 1900212. 10.1002/bies.20200152

Miller D, Hawley R. (2017). Meiotic recombination: taking the path less traveled. Current Biology 27: R19–R41. 10.1016/j.cub.2016.11.002

Nagy B. (2018). *Nothobranchius ditte*, a new species of annual killifish from the Lake Mweru basin in the Democratic Republic of the Congo (Teleostei: Nothobranchiidae). Ichthyo-logical Exploration of Freshwaters 28: 115–134. 10.1002/aqc.3741

Nagy B, Watters BR. (2022). A review of the conservation status of seasonal *Nothobranchius* fishes (Teleostei: Cyprinodontiformes), a genus with a high level of threat, inhabiting ephemeral wetland habitats in Africa. Aquatic Conservation: Marine and Freshwater Ecosystems 32: 199–216. 10.1002/aqc.3741

Nambiar M, Smith GR. (2016). Repression of harmful meiotic recombination in centromeric regions. Seminars in Cell & Developmental Biology 54: 188–197. 10.1016/j.semcdb.2016.01.042

Novák P, Ávila Robledillo L, Koblížková A, Vrbová I, Neumann P, Macas J. (2017). TAREAN: A computational tool for identification and characterization of satellite DNA from unassembled short reads. Nucleic Acids Res 45: e111–e111. 10.1093/nar/gkx257

Olito C, Ponnikas S, Hansson B, Abbott JK. (2022). Consequences of partially recessive deleterious genetic variation for the evolution of inversions suppressing recombination between sex chromosomes. Evolution 76: 1320–1330. 10.1111/evo.14496

Pan Q, Kay T, Depincé A, Adolfi M, Schartl M, Guiguen Y, Herpin A. (2021). Evolution of master sex determiners: TGF-β signalling pathways at regulatory crossroads. Philosophical Transactions of the Royal Society B 376: 20200091. 10.1098/rstb.2020.0091

Peichel CL, McCann SR, Ross JA, Naftaly AFS, Urton JR, Cech JN, Grimwood J, Schmutz J, Myers RM, Kingsley DM, White MA. (2020). Assembly of the threespine stickleback Y chromosome reveals convergent signatures of sex chromosome evolution. Genome Biology 21: 177. 10.1186/s13059-020-02097-x

Pennell MW, Kirkpatrick M, Otto SP, Vamosi JC, Peichel CL, Valenzuela N, Kitano J. (2015). Y Fuse? Sex chromosome fusions in fishes and reptiles. PLOS Genetics 11: e1005237. 10.1371/journal.pgen.1005237

Ponnikas S, Sigeman H, Abbott JK, Hansson B. (2018). Why do sex chromosomes stop recombining? Trends in Genetics 34: 492–503. 10.1016/j.tig.2018.04.001

Quinlan AR, Hall IM. (2010). BEDTools: A flexible suite of utilities for comparing genomic features. Bioinformatics 26: 841–842. 10.1093/bioinformatics/btq033

Ráb P, Roth P. (1988). Cold-blooded vertebrates. In: Balicek P, Forejt J, Rubeš J (eds) Methods of chromosome analysis. Cytogenetická Sekce Československé Biologické Společnosti při CSAV, Brno, Czech Republic, pp 115–124.

Ranallo-Benavidez TR, Jaron KS, Schatz MC. (2020). GenomeScope 2.0 and Smudgeplot for reference-free profiling of polyploid genomes. Nature Communications 11: 1432. 10.1038/s41467-020-14998-3

Reichwald K, Lauber C, Nanda I, Kirschner J, Hartmann N, Schories S, Gausmann U, Taudien S, Schilhabel MB, Szafranski K, Glöckner G, et al. (2009). High tandem repeat content in the genome of the short-lived annual fish *Nothobranchius furzeri*: A new vertebrate model for aging research. Genome Biology 10: R16. 10.1186/gb-2009-10-2-r16

Reichwald K, Petzold A, Koch P, Downie BR, Hartmann N, Pietsch S, Baumgart M, Chalopin D, Felder M, Bens M, et al. (2015). Insights into sex chromosome evolution and aging from the genome of a short-lived fish. Cell 163: 1527–1538. 10.1016/j.cell.2015.10.071

Renner S, Li Y, Wang D, Sun P, Zhao J, Shan L, Ru D, Ren G, Ma T, Liu J. (2023). Evolution of heteromorphic XY chromosomes in sea buckthorn via chromosomal fusion followed by inversions and tissue-specific dosage compensation. Preprint (Research Square) 10.21203/rs.3.rs-3264004/v1

Richter A, Mörl H, Thielemann M, Kleemann M, Geißen R, Schwarz R, Albertz C, Koch P, Petzold A, Groth M, Hartmann N, Herpin A, Englert C. (2023). The Tgf-β family member Gdf6Y determines the male sex in *Nothobranchius furzeri* by suppressing oogenesis-inducing genes. Preprint (bioRxiv), 2023–05. 10.1101/2023.05.26.542338

Sahara K, Marec F, Traut W. (1999). TTAGG telomeric repeats in chromosomes of some insects and other arthropods. Chromosome Research 7: 449–460. 10.1023/A:1009297729547

Sardell JM, Josephson MP, Dalziel AC, Peichel CL, Kirkpatrick M. (2021). Heterogeneous histories of recombination suppression on stickleback sex chromosomes. Molecular biology and Evolution 38: 4403–4418. 10.1093/molbev/msab179

Sardell JM, Kirkpatrick M. (2020). Sex differences in the recombination landscape. The American Naturalist 195: 361–379. 10.1086/704943

Saunders PA, Neuenschwander S, Perrin N. (2018). Sex chromosome turnovers and genetic drift: a simulation study. Journal of Evolutionary Biology 31: 1413–1419. 10.1111/jeb.13336

Saunders PA. (2019). Sex chromosome turnovers in evolution. eLS 10.1002/9780470015902.a0028747

Sember A, Bohlen J, Šlechtová V, Altmanová M, Symonová R, Ráb P. (2015). Karyotype differentiation in 19 species of river loach fishes (Nemacheilidae, Teleostei): Extensive variability associated with rDNA and heterochromatin distribution and its phylogenetic and ecological interpretation. BMC Evolutionary Biology 15: 251. 10.1186/s12862-015-0532-9

Sember A, Nguyen P, Perez MF, Altmanová M, Ráb P, Cioffi MB. (2021). Multiple sex chromosomes in teleost fishes from a cytogenetic perspective: State of the art and future challenges. Philosophical Transactions of the Royal Society B 376: 20200098. 10.1098/rstb.2020.0098

Schartl M, Schmid M, Nanda I. (2016). Dynamics of vertebrate sex chromosome evolution: From equal size to giants and dwarfs. Chromosoma 125: 553–571. 10.1007/s00412-015-0569-y

Smit AF, Hubley R. (2008–2015). RepeatModeler-1.0. Retrieved from http://www.repeatmasker.org

Smit AFA, Hubley R, Green P. (2013–2015). RepeatMasker Open-4.0. Retrieved from http://www.repeatmasker.org. Accessed 2 Jan 2022

Smith SH, Hsiung K, Böhne A. (2023). Evaluating the role of sexual antagonism in the evolution of sex chromosomes: new data from fish. Current Opinion in Genetics & Development 81: 102078. 10.1016/j.gde.2023.102078

Stanke M, Morgenstern B. (2005). AUGUSTUS: A web server for gene prediction in eukaryotes that allows user-defined constraints. Nucleic Acids Research 33: W465– W467. 10.1093/nar/gki458

Štundlová J, Hospodářská M, Lukšíková K, Voleníková A, Pavlica T, Altmanová M, Richter A, Reichard M, Dalíková M, Pelikánová Š, Marta A, Simanovsky SA, Hiřman M, Jankásek M, Dvořák T, Bohlen J, Ráb P, Englert C, Nguyen P, Sember A. (2022). Sex chromosome differentiation via changes in the Y chromosome repeat landscape in African annual killifishes *Nothobranchius furzeri* and *N. kadleci*. Chromosome Research 30: 309–333. 10.1007/s10577-022-09707-3

van der Merwe PDW, Cotterill FPD, Kandziora M, Watters BR, Nagy B, Genade T, Flügel TJ, Svendsen DS, Bellstedt DU. (2021). Genomic fingerprints of palaeogeographic history: The tempo and mode of rift tectonics across tropical Africa has shaped the diversification of the killifish genus *Nothobranchius* (Teleostei: Cyprinodontiformes). Molecular Phylogenetics and Evolution 158: 106988. 10.1016/j.ympev.2020.106988

van Doorn GS, Kirkpatrick M. (2007). Turnover of sex chromosomes induced by sexual conflict. Nature 449:909–912. 10.1038/nature06178

Vara C, Paytuví-Gallart A, Cuartero Y, Álvarez-González L, Marín-Gual L, Garcia F, Florit-Sabater B, Capilla L, Sanchéz-Guillén RA, Sarrate Z, Cigliano RA, Sanseverino W, Searle JB, Ventura J, Marti-Renom MA, Le Dily F, Ruiz-Herrera A. (2021). The impact of chromosomal fusions on 3D genome folding and recombination in the germ line. Nature Communications 12: 2981. 10.1038/s41467-021-23270-1

Veller C, Muralidhar P, Constable GWA, Nowak MA. (2017). Drift-induced selection between male and female heterogamety. Genetics 207: 711–727. 10.1534/genetics.117.300151

Vicoso B. (2019). Molecular and evolutionary dynamics of animal sex-chromosome turnover. Nature Ecology & Evolution 3: 1632–1641. 10.1038/s41559-019-1050-8

Vicoso B, Bachtrog D. (2015). Numerous transitions of sex chromosomes in Diptera. PLOS Biology 13: e1002078. 10.1371/journal.pbio.1002078

Voleníková A, Lukšíková K, Mora P, Pavlica T, Altmanová M, Štundlová J, Pelikánová Š, Simanovsky SA, Jankásek M, Reichard M, Nguyen P, Sember A. (2023). Fast satellite DNA evolution in *Nothobranchius* annual killifishes. Chromosome Research 31: 33. 10.1007/s10577-023-09742-8

Volff JN, Nanda I, Schmid M, Schartl M. (2007). Governing sex determination in fish: regulatory putsches and ephemeral dictators. Sexual Development 1: 85–99. 10.1159/000100030

Völker M, Ráb P. (2015). Direct chromosome preparation from regenerating fin tissue. In: Ozouf-Costaz C, Pisano E, Foresti F, and de Almeida-Toledo LF (eds) Fish cytogenetic techniques: ray-fin fishes and chondrichthyans. CRC Press, Inc, Endfield, pp 37–41. 10.1201/b18534-4

Vrtílek M, Žák J, Pšenička M, & Reichard M. (2018). Extremely rapid maturation of a wild African annual fish. Current Biology 28: R822–R824. 10.1016/j.cub.2018.06.031

Wen M, Pan Q, Jouanno E, Montfort J, Zahm M, Cabau C, Klopp C, Iampietro C, Roques C, Bouchez O, et al. (2022). An ancient truncated duplication of the anti-Müllerian hormone receptor type 2 gene is a potential conserved master sex determinant in the Pangasiidae catfish family. Molecular Ecology Resources. 22: 2411–2428. 10.1111/1755-0998.13620

Wildekamp RH. (1996). A world of killies. Atlas of the oviparous cyprinodontiform fishes of the world (Vol. III). American Killifish Association, Mishawaka, 330 pp.

Wildekamp RH. (2004). A world of killies – atlas of the oviparous cyprinodontiform fishes of the world (Vol. IV). The American Killifish Association, Elyria, Ohio, 398 pp.

Willemsen D, Cui R, Reichard M, Valenzano DR. (2020). Intra-species differences in population size shape life history and genome evolution. eLife 9: e55794. 10.7554/eLife.55794

Wright AE, Dean R, Zimmer F, Mank JE. (2016). How to make a sex chromosome. Nature Communications 7: 12087. 10.1038/ncomms12087

Xiao Y, Xiao Z, Ma D, Zhao C, Liu L, Wu H, et al. (2020) Chromosome-level genome reveals the origin of neo-Y chromosome in the male barred knifejaw *Oplegnathus fasciatus*. iScience 23:101039. 10.1016/j.isci.2020.101039

Xu D, Sember A, Zhu Q, de Oliveira EA, Liehr T, Al-Rikabi ABH, Xiao Z, Song H, Cioffi Cioffi MB. (2019). Deciphering the origin and evolution of the X_1_X_2_Y system in two closely-related *Oplegnathus* species (Oplegnathidae and Centrarchiformes). International Journal of Molecular Sciences, 2 : 3571. 10.3390/ijms20143571

Yang F, Trifonov V, Ng BL, Kosyakova N, Carter NP. (2009). In: Liehr T (ed) Generation of paint probes by flowsorted and microdissected chromosomes. Fluorescence in situ hybridization (FISH)—application guide, 2nd edn. Springer, Berlin, pp 35–52

Yano CF, Bertollo LAC, Ezaz T, Trifonov V, Sember A, Liehr T, Cioffi MB. (2017). Highly conserved Z and molecularly diverged W chromosomes in the fish genus *Triportheus* (Characiformes, Triportheidae). Heredity 118: 276–283. 10.1038/ hdy.2016.83F

Yoshida K, Rödelsperger C, Röseler W, Riebesell M, Sun S, Kikuchi T, Sommer RJ. (2023). Chromosome fusions repatterned recombination rate and facilitated reproductive isolation during *Pristionchus* nematode speciation. Nature Ecology & Evolution 7: 424–439. 10.1038/s41559-022-01980-z

Yoshido A, Sahara K, Yasukochi Y. Silk moths (Lepidoptera) (2015) In: Sharakhov IV (ed) Protocols for cytogenetic mapping of arthropod genomes. CRC Press, Boca Raton, FL, USA, pp 219–256

Zhou Q, Zhang J, Bachtrog D, An N, Huang Q, Jarvis ED, Gilbert MTP, Zhang G. (2014). Complex evolutionary trajectories of sex chromosomes across bird taxa. Science 346: 1246338. 10.1126/science.1246338

